# BIONIC: Biological Network Integration using Convolutions

**DOI:** 10.1101/2021.03.15.435515

**Authors:** Duncan T. Forster, Sheena C. Li, Yoko Yashiroda, Mami Yoshimura, Zhijian Li, Luis Alberto Vega Isuhuaylas, Kaori Itto-Nakama, Daisuke Yamanaka, Yoshikazu Ohya, Hiroyuki Osada, Bo Wang, Gary D. Bader, Charles Boone

**Author notes:** Equal contribution (Duncan T. Forster, Sheena C. Li).

## Abstract

Biological networks constructed from varied data, including protein-protein interactions, gene expression data, and genetic interactions can be used to map cellular function, but each data type has individual limitations such as bias and incompleteness. Network integration promises to address these limitations by combining and automatically weighting input information to obtain a more accurate and comprehensive representation of the underlying biology. However, existing network integration methods may fail to adequately scale to the number of nodes and networks present in genome-scale data, may perform poorly, and may not handle partial network overlap. To address these issues, we developed a deep learning-based network integration algorithm that incorporates a graph convolutional network (GCN) framework to effectively learn dependencies between any input network. Our method, BIONIC (**Bio**logical **N**etwork **I**ntegration using **C**onvolutions), learns features which contain substantially more functional information compared to existing approaches, linking genes that share diverse functional relationships, including co-complex and shared bioprocess annotation. BIONIC can integrate networks in a fully unsupervised manner if functional gene annotations are not available, and it can also leverage available annotations in a semi-supervised manner. BIONIC is scalable in both size and quantity of the input networks, making it feasible to integrate numerous networks on the scale of the human genome. To demonstrate the utility of BIONIC in identifying novel biology, we predicted essential gene chemical-genetic interactions from a small set of diagnostic non-essential gene profiles in yeast, and experimentally validated these predictions. BIONIC correctly predicted many chemical-genetic interactions, and it correctly predicted genes that are required for proper β-1,6-glucan synthesis as significant interactions with the bioactive compound pseudojervine.

## Introduction

High-throughput functional genomics projects produce massive amounts of biological data for thousands of genes. The results of these experiments can be represented as gene-gene interaction networks, which link genes or proteins of related function^1^. For example, protein-protein interactions describe transient or stable physical binding events between proteins^2–7^. Gene co-expression profiles identify genes that share similar patterns of gene expression across multiple experimental conditions, revealing co-regulatory relationships between genes^8,9^. Genetic interactions (e.g. synthetic lethality) link genes that produce an unexpected phenotype when perturbed simultaneously, capturing functional dependencies between genes^10,11^. Each of these data represent functional connections between genes and have varying rates of false-positives and negatives. Data integration has the potential to generate more accurate and more complete functional networks. However, the diversity of experimental methods and results makes unifying and collectively interpreting this information a major challenge.

A number of methods for network integration have been developed with a range of both advantages and disadvantages. For example, many integration algorithms produce networks that retain only global topological features of the original networks, which can be at the expense of important local relationships^12–15^, whereas others fail to effectively integrate networks with partially disjoint node sets^16,17^. Some methods incorporate too much noise in their output, for instance by using more dimensions than necessary to represent their output, which can be detrimental to gene function and functional interaction prediction quality^12–16^. Most data integration approaches do not scale in the number of networks or in the size of the networks required for real world settings^14,16,18^. Supervised methods have traditionally been the most common network integration approach^15,18–20^ and, while highly successful, they require labelled training data to optimize their predictions of known gene functions, which may not be available. Moreover, annotations can be biased and limited, working only with known functional descriptions and reinforcing the existing understanding of gene relationships rather than identifying new ones.

Few biological networks are comprehensive in terms of genome coverage and the overlap in the set of genes captured by different networks is often limited. Some methods ignore this problem by simply integrating genes that are common to all networks^16^, resulting in progressively smaller gene sets as more networks are added, whereas others produce gene features that are dependent on whether a gene is present in all or only some of the input networks^17^. Integration methods generally require each input network to have the same set of genes, so to produce an integrated result that encompasses genes present across all networks (i.e. the union of genes) each network must be extended with any missing genes^12,15–17,21^. However, existing methods often do not distinguish between genes that have zero interactions due to this extension, and genes with zero measured interactions in the original data^12,15–17^.

Unsupervised methods have more recently been explored to perform biological network integration. They automatically identify network structure, such as modules, shared across independent input data and can function in an unbiased manner, using techniques such as matrix factorization^12–14^, cross-diffusion^16^, low-dimensional diffusion state approximation^17^ and multimodal autoencoding^21^. Theoretically, unsupervised network integration methods can provide a number of desirable features such as automatically retaining high-quality gene relationships and removing spurious ones, inferring new relationships based on the shared topological features of many networks in aggregate, and outputting comprehensive results that cover the entire space of information associated with the input data, all while remaining agnostic to any particular view of biological function.

Recently, new methods have been developed that focus on learning compact features over networks^22,23^. These strategies aim to capture the global topological roles of nodes (i.e. genes or proteins) and reduce false positive relationships by compressing network-based node features to retain only the most salient information. However, this approach produces general purpose node features that are not necessarily optimal for the task of interest. Advances in deep learning have addressed this shortcoming with the development of the graph convolutional network (GCN), a general class of neural network architectures which are capable of learning features over networks^24–27^. GCNs can learn compact, denoised node features that are trainable in a network-specific fashion. Additionally, the modular nature of the GCN enables the easy addition of specialized neural network architectures to accomplish a task of interest, such as network integration, while remaining scalable to large input datasets. Compared to general-purpose node feature learning approaches^22,23^, GCNs have demonstrated substantially improved performance for a range of general network tasks, which is a direct result of their effective feature learning capabilities^24,27^. These promising developments motivate the use of the GCN for gene and protein feature learning on biological networks, which can be large and feature highly variable network topologies.

Here we present a general, scalable deep learning framework for network integration called BIONIC (**Bio**logical **N**etwork **I**ntegration using **C**onvolutions) which uses GCNs to learn gene features given many different input networks. BIONIC addresses the aforementioned limitations of existing integration methods, it handles diverse biological networks with only partially overlapping gene sets, and it produces integration results which accurately reflect the underlying network topologies and capture functional information. Moreover, BIONIC is scalable in the number of networks and the size of these networks, and it can perform integration in both unsupervised and semi-supervised scenarios depending on the available data. To demonstrate the utility of BIONIC, we integrate three diverse, high-quality gene and protein interaction networks, to obtain integrated gene features that we compare to a range of function prediction benchmarks. We analyze our findings in the context of those obtained from a wide range of integration methodologies^12,17^, and we show that BIONIC features perform well at both capturing functional information and scaling in terms of the number of networks and network size, while maintaining gene feature quality. Finally, we applied BIONIC network integration towards the analysis of chemical-genetic interactions^28^, which allowed us to make novel predictions about the cellular targets of previously uncharacterized bioactive compounds.

## Results

### BIONIC architecture

BIONIC uses the GCN neural network architecture to learn optimal gene (protein) interaction network features individually, and combines these features into a single, unified representation for each gene (**Fig. 1**). First, the input data, if not already in a network format, are converted to networks (e.g. by gene expression profile correlation) (**Fig. 1a**). Each input network is then run through a sequence of GCN layers (**Fig. 1b**) to produce network-specific gene features. The number of GCN layers used (three layers in our experiments - see **Methods, Supplementary Data File 1**) determines the size of the neighborhood (i.e. genes directly connected to a given gene) used to update the gene features^24^, where one layer would use only the gene’s immediate neighbors, two layers would use the second order neighborhood, and so on. Residual connections are added from the output of each network-specific GCN layer in the sequence to the output of the final GCN in the sequence (**Fig. S1**). This allows BIONIC to learn gene features based on multiple neighborhood sizes rather than just the final neighborhood, while additionally improving training by preventing vanishing gradients^29^. The network-specific features are then summed through a stochastic gene dropout procedure to produce unified gene features which can be used in downstream tasks, such as functional module detection or gene function prediction. To optimize the functional information encoded in its integrated features, BIONIC must have relevant training objectives that facilitate capturing salient features across multiple networks. Here, BIONIC uses an unsupervised training objective, and if some genes have functional labels (such as complex, pathway or bioprocess membership annotations), BIONIC can also use these labels to update its learned features though a semi-supervised objective.

**Figure 1.**
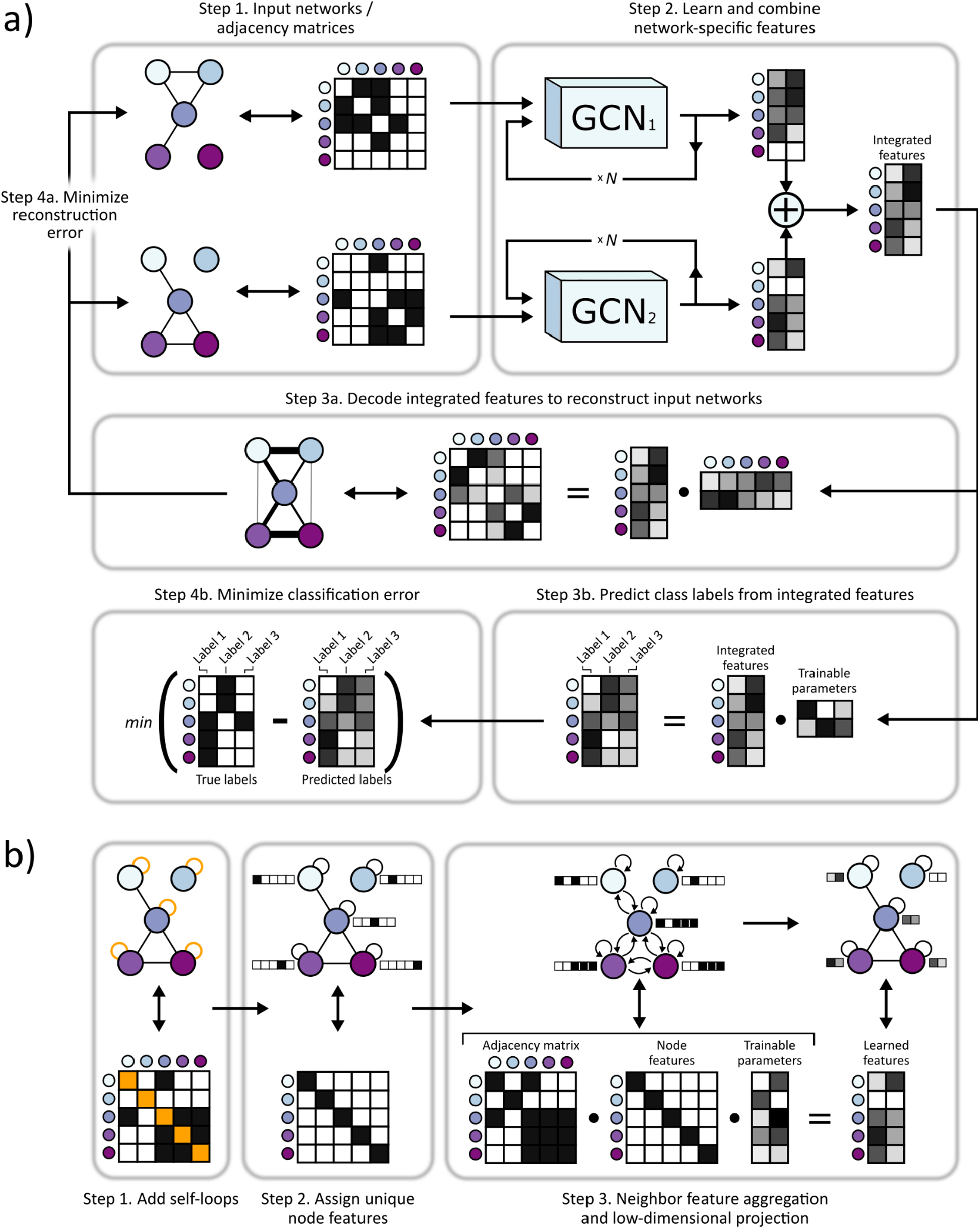
**a)** BIONIC integrates networks as follows: Step 1. Gene interaction networks input into BIONIC are represented as adjacency matrices. Step 2. Each network is passed through a graph convolution network (GCN) to produce network-specific gene features which are then combined into an integrated feature set which can be used for downstream tasks such as functional module detection. The GCNs can be stacked multiple times (denoted by N) to generate gene features encompassing larger neighborhoods. Step 3a. (Unsupervised) BIONIC attempts to reconstruct the input networks by decoding the integrated features through a dot product operation. Step 4a. (Unsupervised) BIONIC trains by updating its weights to reproduce the input networks as accurately as possible. Step 3b. (Semi-supervised) If labelled data is available, BIONIC predicts functional labels for each gene using the learned gene features. Step 4b. (Semi-supervised) BIONIC trains by updating its weights to predict the ground-truth labels and minimize classification error. **b)** The GCN architecture functions by: Step 1. adding self-loops to each network node, Step 2. assigning a “one-hot” feature vector to each node in order for the GCN to uniquely identify the nodes and Step 3. propagating node features along edges followed by a low-dimensional, learned projection to obtain updated node features which encode the network topology.

For the unsupervised objective, BIONIC uses an autoencoder design and reconstructs each input network by mapping the integrated gene features to a network representation (decoding) and minimizing the difference between this reconstruction and the original input networks. By optimizing the fidelity of the network reconstruction, BIONIC forces the learned gene features to encode as much salient topological information present in the input networks as possible, which reduces the amount of spurious information encoded. Indeed, for a set of three yeast networks^5,9,11^, inputting these networks into BIONIC individually tends to produce features with higher performance on several benchmarks compared to the original network format (**Fig. S2**). Biological networks generally overlap only partially in their gene sets. If a gene is missing from a given network, it is not appropriate to reconstruct this gene in the given network, since the adjacency profile of this gene is unknown (i.e. not measured in the experiment). To handle this partial overlap, BIONIC implements a masking procedure which prevents penalizing the reconstruction fidelity of gene interaction profiles in networks where the genes were not originally present (see **Methods**).

For the semi-supervised objective, BIONIC predicts labels for each gene using the integrated gene features and then updates its weights by minimizing the difference between the predictions and a set of user-specified ground-truth functional labels. Here, BIONIC performs multi-label classification, where a given gene may be assigned more than one class label. BIONIC ignores the classification error for any genes lacking ground-truth labels, and so is able to incorporate as much (or as little) labelled information as is available. The semi-supervised classification objective is used in conjunction with the unsupervised network reconstruction objective when gene labels are available, and the unsupervised objective is used on its own when no gene labels are available.

### Evaluation criteria

For the following analyses, we assessed the quality of the input networks and network integration method outputs using three evaluation criteria: (1) gene co-annotation prediction; (2) gene module detection; (3) supervised gene function prediction. First, we used an established precision-recall evaluation strategy^11,30^ to determine how well gene-gene relationships produced by the given method overlapped with gene pairs co-annotated to the same term in a particular functional standard. Second, we evaluated the capacity of each method to produce biological modules by comparing clusters computed from the output of each method to known modules such as protein complexes, pathways, and biological processes. These two evaluations measure the intrinsic quality of the outputs generated by the integration methods, i.e. without training any additional models on top of the outputs. Finally, the supervised gene function prediction evaluation determines how discriminative the method outputs are for predicting known gene functions. Here, a portion of the genes and corresponding labels (known functional classes such as protein complex membership) were held out and used to evaluate the accuracy of a support vector machine classifier^31^, which is trained on the remaining gene features, output from the given integration method, to predict the held-out labels^17^. This constitutes an extrinsic evaluation, indicating how effectively the method outputs can be used in conjunction with an additional classification model.

In the following experiments, to ensure a fair choice of hyperparameters across BIONIC and the integration methods we compared to, we performed a hyperparameter optimization step using an independent set of *Schizosaccharomyces pombe* networks as inputs^32–34^ and a set of Gene Ontology curated pombe protein complexes^35^ for evaluation. The best performing hyperparameters for each approach were used (see **Methods**).

### Evaluation of BIONIC features and input networks

We first used BIONIC to integrate three diverse yeast networks: a comprehensive network of correlated genetic interaction profiles (4,529 genes, 33,056 interactions)^11^, a co-expression network derived from transcript profiles of yeast strains carrying deletions of transcription factors (1,101 genes, 14,826 interactions)^9^, and a protein-protein interaction network obtained from an affinity-purification mass-spectrometry assay (2,674 genes, 7,075 interactions)^5^, which combine for a total of 5,232 unique genes and 53,351 unique interactions (**Fig. 2**, **Supplementary Data File 2**). Compared to the input networks, BIONIC integrated features have equivalent or superior performance on all evaluation criteria over three different functional benchmarks: IntAct protein complexes^36^, Kyoto Encyclopedia of Genes and Genomes (KEGG) pathways^37^ and Gene Ontology biological processes (GO)^35^ (**Fig. 2a, Supplementary Data File 3**). As an additional test, BIONIC produces high-quality features that accurately predict a diverse set of yeast biological process annotations per gene^11^ (**Fig. 2b**). Some categories in this latter test do better than others. These performance patterns were mirrored in the individual input networks, indicating that this is the result of data quality, rather than method bias. For example, BIONIC performance on this per-biological process benchmark is driven primarily by the genetic interaction network, which captures broad biological process gene relationships better than the protein-protein interaction or co-expression networks. We conclude that, BIONIC captures high-quality functional information across diverse input networks, network topologies and gene function categories, and that the BIONIC output features can be used to accurately identify pairwise gene co-annotation relationships, functional modules, and predict gene function.

**Figure 2.**
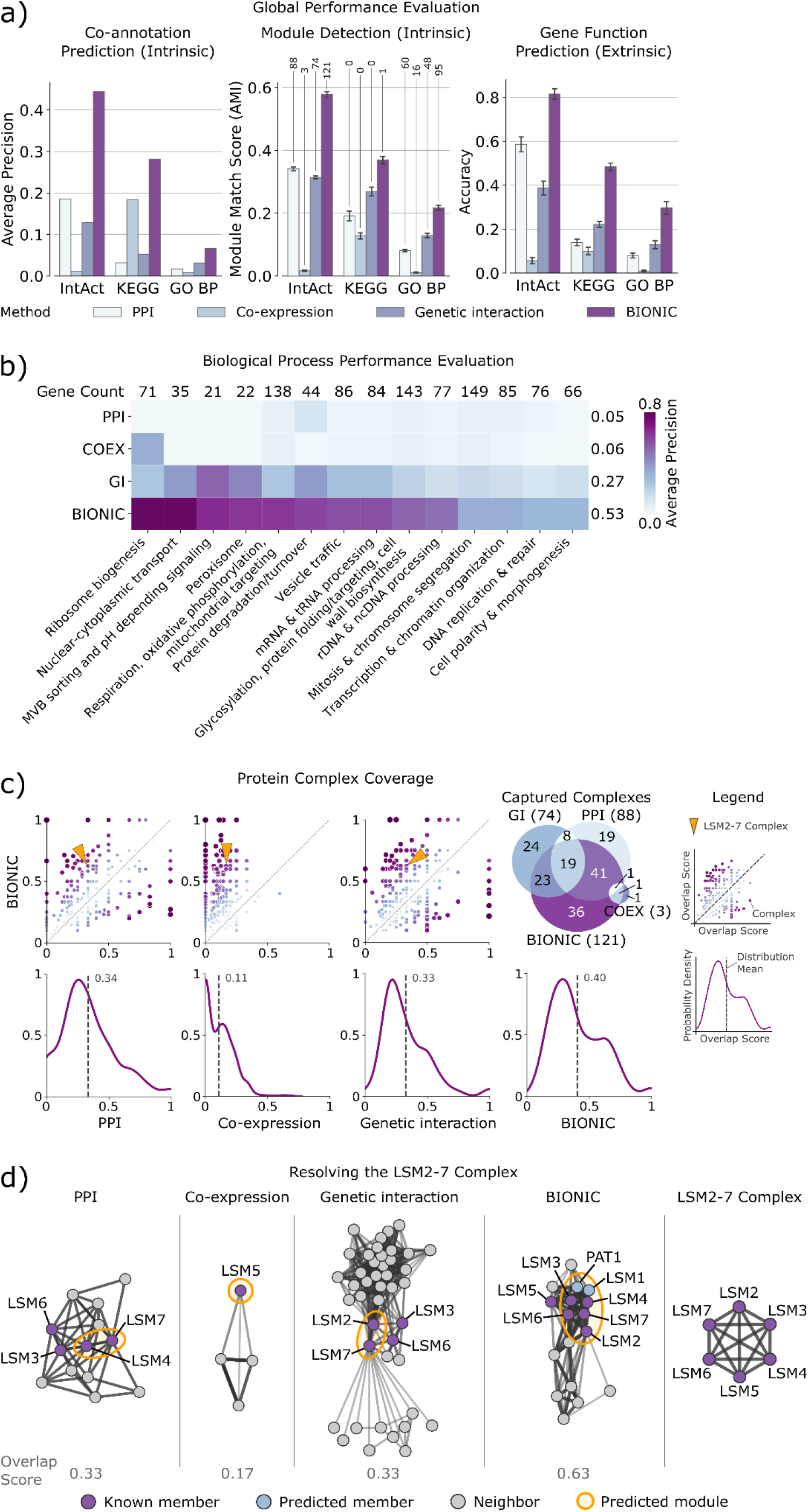
**a)** Co-annotation prediction, module detection, and gene function prediction evaluations for three yeast networks, and BIONIC features from the integration of these networks. The co-annotation and module detection standards contain between 1786 and 4170 genes overlapping the integration results. The module detection standards define between 107 and 1809 modules. The IntAct, KEGG and GO BP gene function prediction standards cover 567, 1770 and 1211 genes overlapping the integration results, and 48, 53 and 63 functional classes, respectively (see **Supplementary Data File 2**). Error bars indicate the 95% confidence interval. Numbers above the module detection bars indicate the number of captured modules, as determined by a 0.5 overlap (Jaccard) score cutoff. **b)** Evaluation of networks and integrated features using high-level functional categories, split by category. Each category contains between 21 and 149 genes overlapping the integration results (denoted by counts above the heatmap columns, see **Supplementary Data File 2**) and the average performance of each method across categories is reported (scores to the right of each row). **c) Top row**: Comparison of overlap scores between known complexes and predicted modules, between BIONIC and the input networks. Each point is a protein complex. The x and y axes indicate the overlap (Jaccard) score, where a value of 0 indicates no members of the complex were captured, and 1 indicates the complex was captured perfectly. The diagonal indicates complexes where BIONIC and the given input network have the same score. Points above the diagonal are complexes where BIONIC outperforms the given network, and points below the diagonal are complexes where BIONIC underperforms the network. The arrows indicate the LSM2-7 complex, shown in d). A Venn diagram describes the overlap of captured complexes (defined as a complex with an overlap score of 0.5 or higher) between the input networks and BIONIC integration. Numbers in brackets denote the total number of captured complexes for the corresponding method. **Bottom row**: The distribution of overlap scores between predicted and known complexes for each network and BIONIC. The dashed line indicates the distribution mean. **d)** Functional relationships between predicted LSM2-7 complex members and genes in the local neighborhood, as given by the three input networks and corresponding BIONIC integration of these networks. The predicted cluster best matching the LSM2-7 complex in each network, based on the module detection analysis in a), is circled. The overlap score of the predicted module with the LSM2-7 complex is shown. Edges correspond to protein-protein interactions in PPI^5^, Pearson correlation between gene profiles in Co-expression^9^ and Genetic Interaction^11^ networks, and cosine similarity between gene features in the BIONIC integration. The complete LSM2-7 complex is depicted on the right. Edge weight corresponds to the strength of the functional relationship (correlation), where a heavier edge implies a stronger functional connection. PPI = Protein-protein interaction, COEX = Co-expression, GI = Genetic interaction, GO = Gene Ontology, BP = Biological process.

We observed that features obtained through BIONIC network integration often outperformed the individual input networks at capturing functional modules (**Fig. 2a**) and captured more modules (**Fig. 2c, Supplementary Data File 4**), demonstrating the utility of the combined features over individual networks for downstream applications such as module detection. Here we treated the network adjacency profiles (rows in the adjacency matrix) as gene features. We then examined how effectively the input networks and integrated BIONIC features captured known protein complexes, by matching each individual known complex to its best matching predicted module and quantifying the overlap (**Fig. 2c**). We then compared the overlap scores from each network to the BIONIC overlap scores to identify complexes where BIONIC performs either better or worse than the input networks. Of 344 protein complexes tested, BIONIC strictly improved 196, 309, 222 complex predictions and strictly worsened 82, 17, 98 complex predictions compared to the input protein-protein interaction, co-expression, and genetic interaction networks, respectively. The distributions of complex overlap scores for each dataset indicate that BIONIC predicts protein complexes more accurately than the input networks on average. Indeed, if we use an overlap score of 0.5 or greater to indicate a successfully captured complex, the integrated BIONIC features, containing information from three networks, capture 121 complexes, compared to 88, 3 and 74 complexes for the individual protein-protein interaction, co-expression, and genetic interaction networks, respectively (**Fig. 2c**). We also repeated this module analysis while optimizing the clustering parameters on a per-module basis, an approach that tests how well each network and BIONIC perform at capturing modules under optimal clustering conditions for each module. Here too, the integrated BIONIC features capture more modules and with a greater average overlap score than the individual input networks (**Fig. S3, S4, Supplementary Data File 5**).

To better understand how BIONIC is able to improve functional gene module detection compared to the input networks, we examined the *LSM2-7* complex, which was identified in our module detection evaluation (**Fig. 2a**) as an example to show how BIONIC effectively combines gene-gene relationships across different networks and recapitulates known biology. The *LSM2-7* complex localizes to the yeast nucleoli and is involved in the biogenesis or function of the small nucleolar RNA snR5^38^. *LSM2-7* is made up of the protein products of six genes - *LSM2*, *LSM3*, *LSM4*, *LSM5*, *LSM6* and *LSM7*. We found that the cluster which best matched the *LSM2-7* complex in each input network only captures a subset of the full complex (**Supplementary Data File 4**). The BIONIC module, however, contains five out of six members of the *LSM2-7* complex, along with two additional members: *LSM1* and *PAT1*, which are functionally associated with the *LSM2-7* complex^39^. The missing member, *LSM5*, is in the local neighborhood of the cluster in the BIONIC feature space. We examined the best-matching clusters and their local neighborhood, consisting of genes that show a direct interaction with predicted members of the *LSM2-7* complex, in the input networks, and in a profile similarity network obtained from the integrated BIONIC features of these networks (**Fig. 2d**). We found that both the PPI and genetic interaction networks captured two members of the *LSM2-7* complex, with two additional members in the local neighborhood. The co-expression network only identified one complex member, and the local neighborhood of the best matching module did not contain any additional known complex members. Finally, BIONIC utilized the interaction information across input networks to better identify the *LSM2-7* module, with the addition of two functionally related proteins. This analysis demonstrates the utility of BIONIC for identifying meaningful biological modules by effectively combining information across input networks. Indeed, when we optimized the module detection procedure to specifically resolve the *LSM2-7* complex, we found that BIONIC was able to capture the complex with a higher overlap score (0.83) than any of the input networks (0.33, 0.17 and 0.50 for the PPI, co-expression and genetic interactions networks, respectively), and it outperformed other integration methods (0.43, 0.22, 0.44, 0.60 and 0.68 for the Union, iCell, deepNF, Mashup and multi-node2vec methods, respectively) (**Supplementary Data File 5**).

### Evaluation of BIONIC and established unsupervised integration methods

We compared network integration results from BIONIC (**Fig. 2**) to those derived from several different established integration approaches: a naive union of networks (Union), a non-negative matrix tri-factorization approach (iCell)^12^, a deep learning multi-modal autoencoder (deepNF)^21^, a low-dimensional diffusion state approximation approach (Mashup)^17^, and a multi-network extension of the node2vec^23^ model (multi-node2vec)^40^ (**Fig. 3**). These unsupervised integration methods cover a wide range of methodologies and the major possible output types (networks for Union and iCell, features for deepNF, Mashup and multi-node2vec). BIONIC performs as well as, or better than the tested integration methods across all evaluation types and benchmarks (**Fig. 3a**). We also evaluated BIONIC and the other integration approaches on a per-biological process basis (**Fig. 3b**). Here we found BIONIC generally outperforms the established integration approaches on each biological process, with the exception of several biological processes when compared to deepNF. Averaging over the performance for each biological process, we found BIONIC performs on par with deepNF (average precision of 0.53 for BIONIC compared to 0.52 for deepNF). DeepNF performs competitively on the per-biological process evaluations (**Fig. 3b**), but it underperforms on the global performance evaluations (**Fig. 3a**). The per-biological process evaluations assess how well a method predicts large-scale biological process co-annotation, whereas the global performance evaluations measure how well a method predicts smaller-scale functional modules (i.e. protein complexes). This discrepancy in performance indicates deepNF is able to capture broad-scale functional organization, but it fails to resolve smaller functional modules. BIONIC performs well on both of these evaluations, however, indicating it can learn gene features which resolve both broad and detailed functional organization.

**Figure 3.**
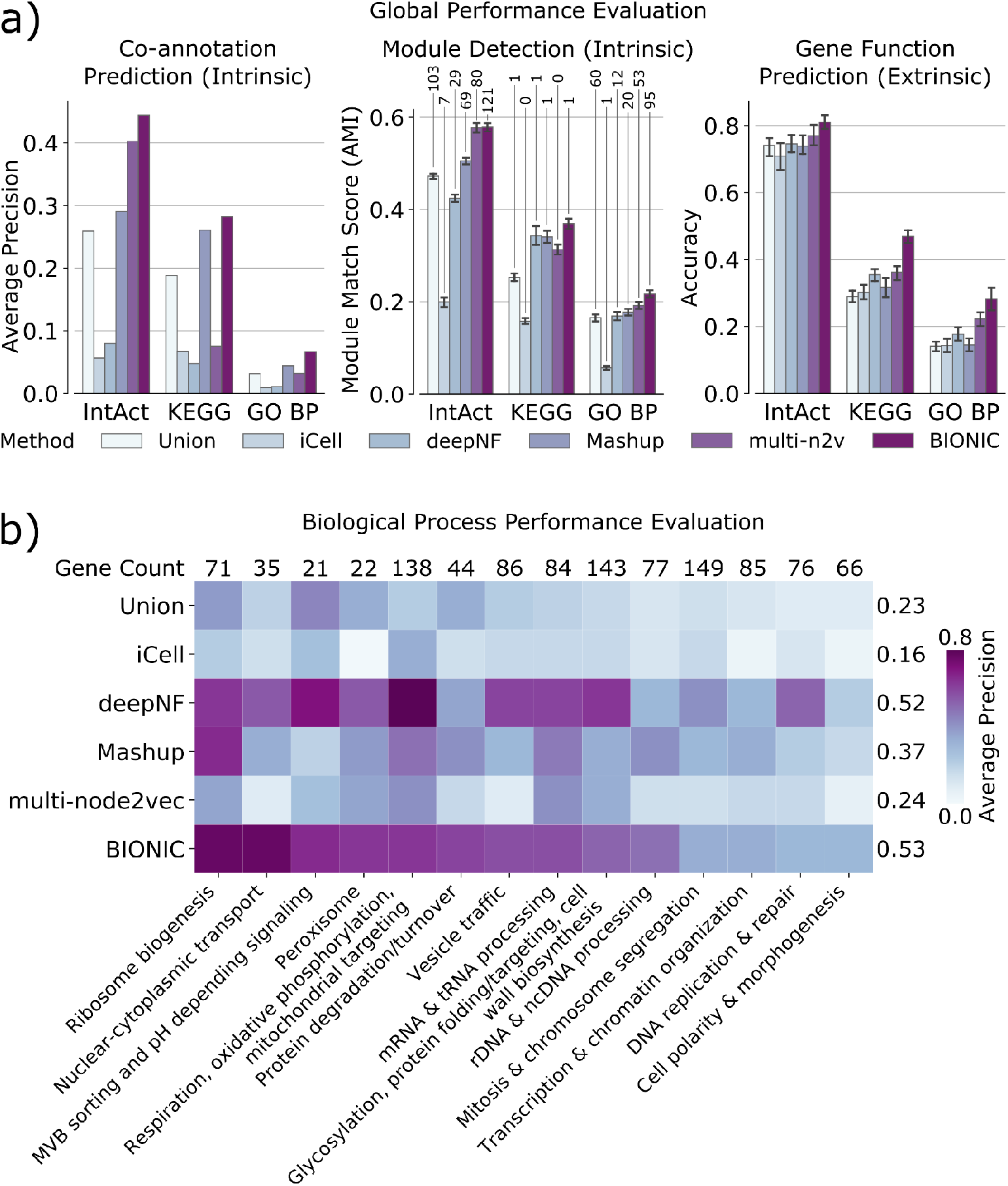
**a)** Co-annotation prediction, module detection, and gene function prediction evaluations for three yeast networks integrated by the tested unsupervised network integration methods. The input networks and evaluation standards are the same as in **Fig. 2**. Error bars indicate the 95% confidence interval. Numbers above the module detection bars indicate the number of captured modules, as determined by a 0.5 overlap (Jaccard) score cutoff. **b)** Evaluation of integrated features using high-level functional categories, split by category. Each category contains between 21 and 149 genes overlapping the integration results (denoted by counts above the heatmap columns, see **Supplementary Data File 2**) and the average performance of each method across categories is reported (scores to the right of each row). PPI = Protein-protein interaction, GO = Gene Ontology, BP = Biological process.

### Evaluation of BIONIC in a semi-supervised setting

We also tested how BIONIC performs in a semi-supervised setting (**Fig. 4**). Here, we compared BIONIC trained with no labelled data (unsupervised), BIONIC trained with a held-out set of functional labels given by IntAct, KEGG, and GO (semi-supervised), and a supervised integration algorithm using the same labels (GeneMANIA^15^). For each of these methods, we integrated the yeast protein-protein interaction, co-expression, and genetic interaction networks from the **Fig. 2** analysis. 20% of genes in each benchmark (IntAct, KEGG, GO) were randomly held out and used as a test set, while the remaining 80% of genes were used for training. The unsupervised BIONIC did not use any gene label information for training, but it was evaluated using the same test set as the supervised methods to ensure a consistent performance comparison. To control for variability in the train-test set partitioning, this procedure was repeated 10 times and the average performance across test sets was reported (see **Methods**). We found that adding labelled data can significantly improve the features BIONIC learns and these features also outperform the integration results produced by the supervised GeneMANIA method. We also found that even without labelled data, BIONIC performs as well as, or exceeds GeneMANIA performance. Notably, the performance of the unsupervised and semi-supervised BIONIC is similar for gene function prediction. This indicates unsupervised BIONIC features are already sufficiently discriminative for classifiers to perform well. Thus, BIONIC can be used effectively in both an unsupervised and semi-supervised setting, which demonstrates its versatility as a biological network integration algorithm.

**Figure 4.**
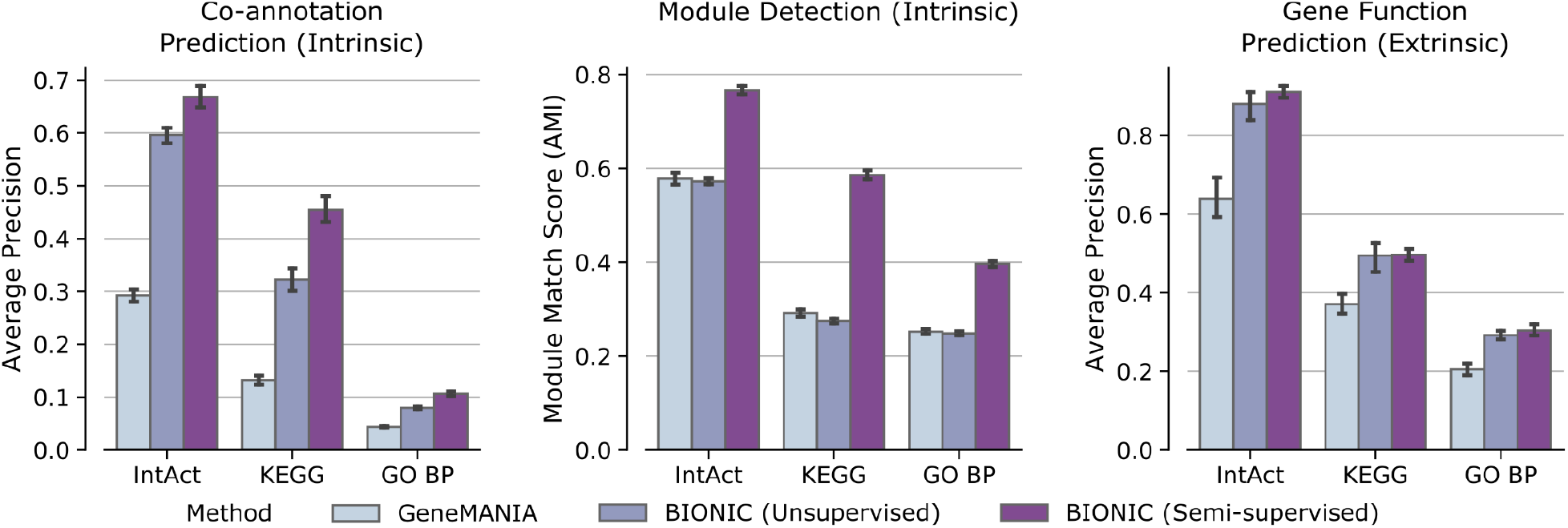
Performance comparison between a supervised network integration algorithm trained with labelled data (GeneMANIA), BIONIC trained without any labelled data (Unsupervised), and BIONIC trained with labelled data (Semi-supervised). Bars indicate the average performance over 10 trials of random train-test splits for the given benchmark (see **Methods**). Error bars indicate the 95% confidence interval. GO = Gene Ontology, BP = Biological process

### Scalability of BIONIC and established integration approaches

High-throughput experiments have led to a rapidly growing wealth of biological networks. For the well studied systems, including yeast and human, there are hundreds of available networks which, when unified, often include close to a genome-wide set of genes. Ideally, all of these networks could be unified to improve available gene function descriptions. However, many unsupervised integration methods either cannot run with many input networks or networks with large numbers of genes, or they scale with reduced performance. To test network input scalability, we randomly sampled progressively larger sets of yeast gene co-expression networks (**Fig. 5a, Supplementary Data File 2**) and assessed the performance of the resulting integrations of these sets. We similarly tested node scalability by randomly subsampling progressively larger gene sets of four human protein-protein interaction networks^3,6,7,41^ (**Fig. 5b, Supplementary Data File 2**). BIONIC can integrate numerous networks (**Fig. 5a**), and networks with many nodes (**Fig. 5b**), outperforming all other methods assessed for progressively more and larger networks. To achieve this scalability, BIONIC takes advantage of the versatile nature of deep learning technology by learning features for small batches of genes and networks at a time, reducing the computational resources required for any specific training step. To learn gene features over large networks, BIONIC learns features for random subsets of genes at each training step, and randomly subsamples the local neighborhoods of these genes to perform the graph convolution (see **Methods**), maintaining a small overall computational footprint. This subsampling allows BIONIC to integrate networks with many genes, whereas methods like Mashup can only do so with an approximate algorithm which substantially reduces integration performance (**Fig. S5**). To integrate many networks, BIONIC uses a network-wise sampling approach, where a random subset of networks is integrated at a time during each training step. This reduces the number of parameter updates required at once, since only GCNs corresponding to the subsampled networks are updated in a given training step.

**Figure 5.**
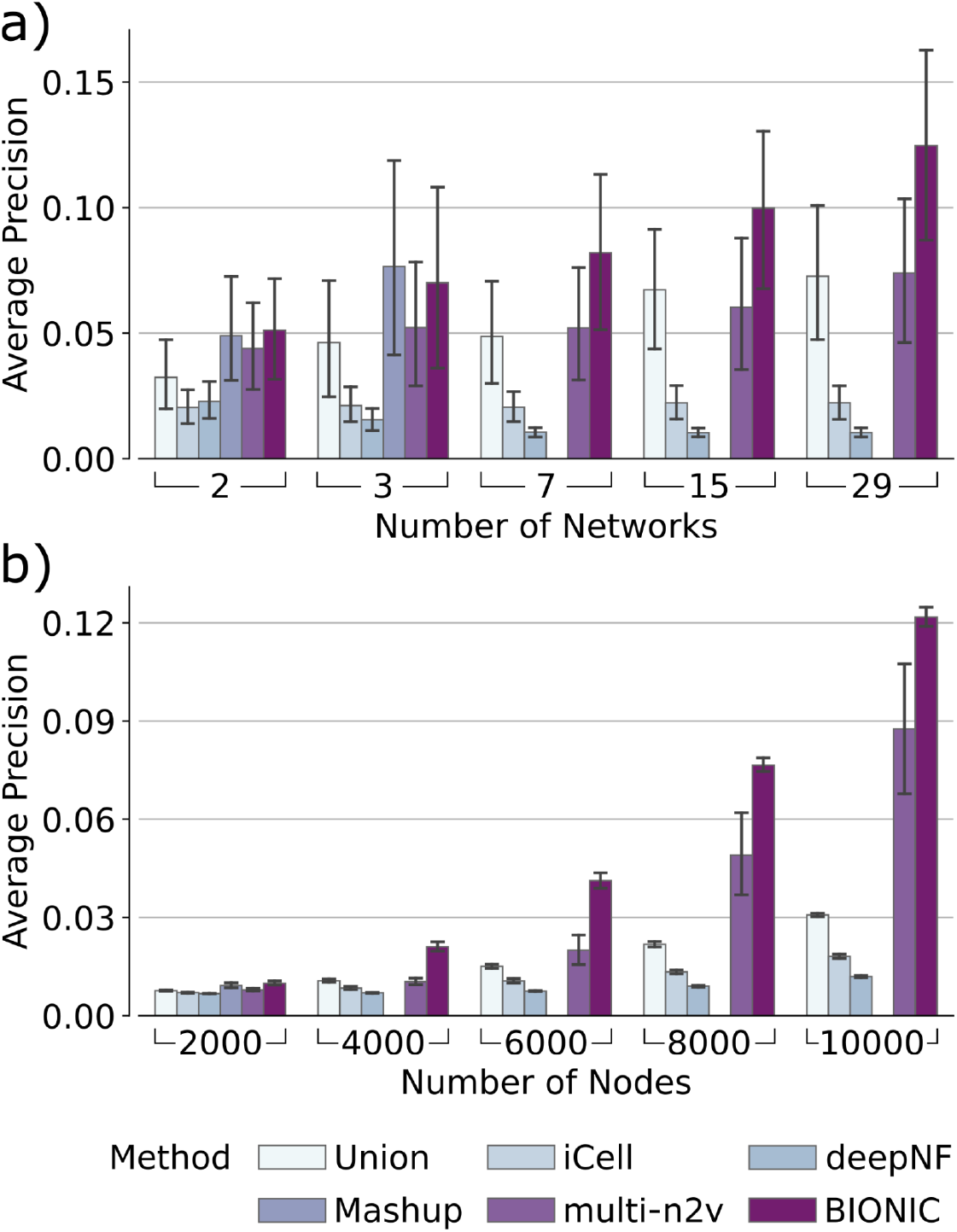
**a)** Performance of integrating various numbers of randomly sampled yeast co-expression input networks on KEGG Pathways gene co-annotations. **b)** Performance of integrating four human protein-protein interaction networks over a range of sub-sampled nodes (genes) on CORUM Complexes protein co-annotations. In these experiments the Mashup method failed to scale to a) 7 or more networks and b) 4000 or more nodes, as indicated by the absence of bars in those cases (see **Methods**). Error bars indicate the 95% confidence interval. multi-n2v = multi-node2vec

### BIONIC predictions of chemical-genetic interactions

We asked if BIONIC can generate new, testable biological hypotheses. Chemical genetic approaches analyze the effects of mutations on cell growth in response to compound treatment, and can be used to systematically predict the molecular targets of uncharacterized compounds^42^. For example, chemical-genetic analysis of heterozygous diploid yeast essential gene deletion mutants has been used to predict compound targets^43^. If a heterozygous diploid mutant carries half the normal copy of a compound’s target genes, it is often specifically hypersensitive to the compound. Similarly, if a conditional temperature sensitive mutant carries a mutation that compromises the activity of a compound’s target gene, it is often specifically hypersensitive to the compound^44,45^.

Previously, we generated a data set of chemical-genetic screens, consisting of a pool of deletion mutants of 289 nonessential genes (diagnostic pool) and 1522 compounds^28^. Using this data, we used BIONIC to predict chemical sensitivities for a wider set of 883 essential genes across a subset of 50 compounds. For the compound selection procedure, we used the BIONIC integrated protein-protein interaction network, co-expression network, and genetic interaction network features from the **Fig. 2** analysis. We selected compounds to study by identifying those that BIONIC predicts well within the diagnostic pool data. We did this by partitioning sensitive genes from each compound into train and test sets, and we used the BIONIC features to predict the test set genes using the training genes as input (see **Methods**). The top 50 compounds, for which sensitive genes were most successfully predicted, were selected for further analysis. Sensitive essential gene predictions for each of the 50 chosen compounds were generated in a similar way to the compound selection procedure, with predictions being made on yeast essential genes rather than the diagnostic pool genes (see **Methods**).

The BIONIC essential gene sensitivity predictions were experimentally validated using profiles for the compound set from a chemical-genetic screen using a collection of temperature sensitive (TS) yeast mutants (**Supplementary Data File 6**). A DNA bar-coded collection of 1181 mutants containing TS alleles spanning 873 genes was constructed in a yeast genetic background that conferred drug hypersensitivity (*pdr1Δpdr3Δsnq2Δ*). The TS mutant collection was pooled and screened against the compound set. Mutant-specific barcodes were amplified from each compound-treated pool, and Illumina sequencing was used to quantify the relative abundance of TS mutant strains in the presence of each compound. Sequencing data was processed using BEAN-counter software to quantify chemical-genetic interactions and eliminate non-specific technical effects^46^. Further statistical analysis was conducted to identify chemical-genetic interactions that satisfied a “far outlier” cut-off (see **Methods**), which were then compared to the sensitive genes predicted by BIONIC.

Out of 156 essential genes experimentally identified as sensitive to the set of 50 screened compounds, BIONIC successfully predicted 35. BIONIC significantly predicts sensitive genes for 13 out of 50 compounds under an ordered Fisher’s exact test. We also assessed more broadly whether BIONIC can correctly predict the biological process a given compound’s sensitive genes are annotated to. BIONIC sensitive gene predictions were statistically enriched (Fisher’s exact test) for 27 out of 62 annotated biological processes across compounds (**Fig. 6a**). We compared the quality of BIONIC’s predictions to a random baseline (**Fig. 6b**). Here, we generated 1000 random permutations of the BIONIC PEG feature gene labels and computed sensitive essential gene predictions for the 50 screened compounds, as described previously. We found BIONIC sensitive gene and bioprocess predictions were substantially more accurate than the random permutations, indicating the BIONIC PEG features encode relevant information for the prediction of chemical-genetic interactions. We looked at the 13 significantly predicted compounds in more detail to see which sensitive gene predictions BIONIC correctly predicted and the corresponding ranks of those genes in the prediction list (**Fig. 6c**). We observed that for 8 out of 13 compounds, the correct BIONIC predictions rank in the top 10 most sensitive interactions. BIONIC predictions and experimental results for the 50 selected compounds can be found in **Supplementary Data File 7**.

**Figure 6.**
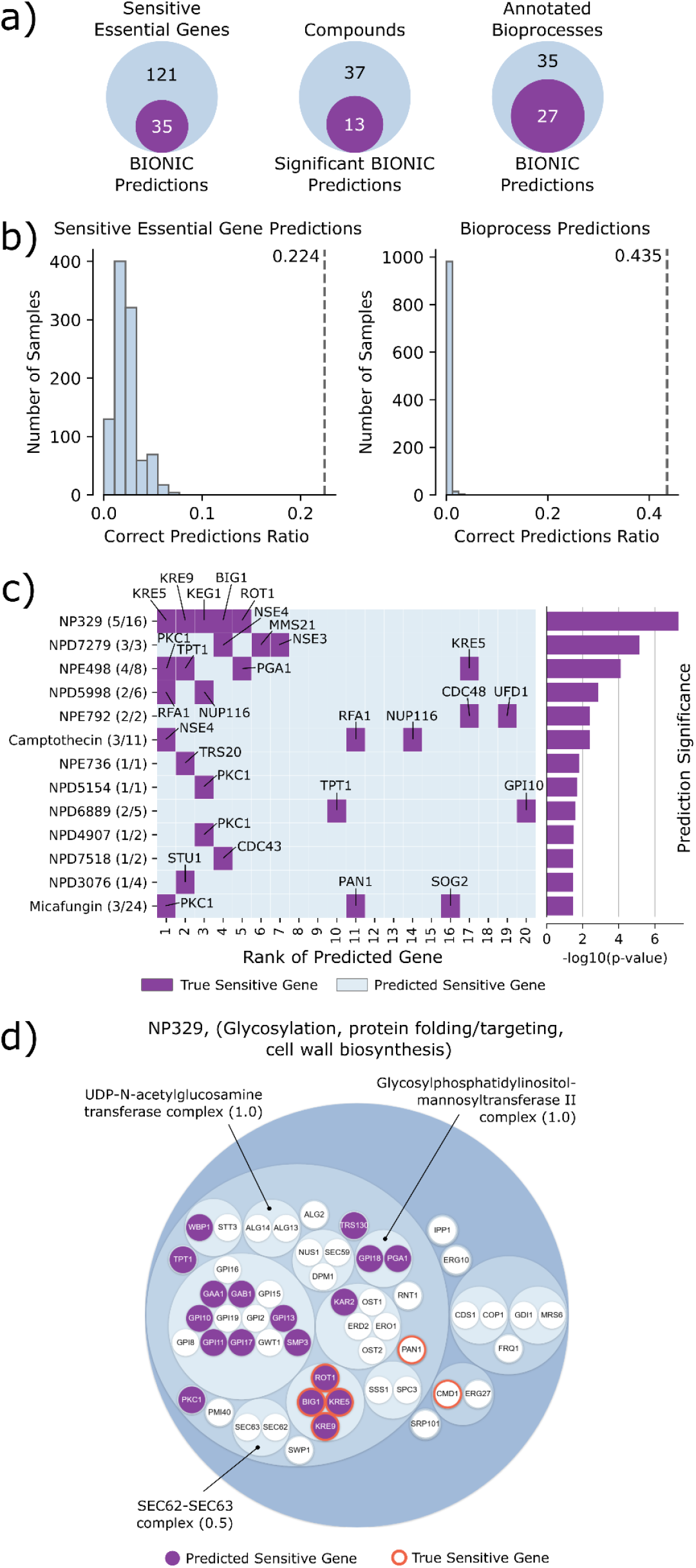
**a)** From left to right: the number of correct BIONIC sensitive essential gene predictions across the 50 screened compounds, the number of compounds BIONIC significantly predicted sensitive essential genes for (ordered Fisher’s exact test), and the number of correctly predicted sensitive essential gene annotated bioprocesses, based on the bioprocess enrichment of BIONIC predictions for each compound. **b)** A comparison of correctly predicted sensitive genes (left) and correctly predicted biological process annotations (right) between BIONIC predictions (dashed line) and 1000 random permutations of BIONIC features gene labels (histogram). Correct prediction ratio is the number of correct predictions divided by the number of total sensitive essential genes (left) or annotated biological processes (right) across the 50 screened compounds. **c)** Rank of BIONIC sensitive essential gene predictions for the 13 significantly predicted compounds. The number of correctly predicted genes out of total sensitive genes are shown in parentheses beside each compound name. The statistical significance of the BIONIC predictions for each compound is displayed in the bar plot on the right. **d)** Hierarchical organization of essential genes in the glycosylation, protein folding/targeting, cell wall biosynthesis bioprocess based on integrated BIONIC features. Smallest circles correspond to genes, larger circles indicate clusters of genes. 6 genes sensitive to the NP329 compound are indicated with orange borders, and corresponding BIONIC predictions lying in the bioprocess are indicated as purple circles. Captured protein complexes in the bioprocess are annotated and the corresponding overlap score (Jaccard) with the true complex is given in parentheses.

We examined the best predicted compound, NP329, in more detail. NP329 is a pseudojervine from the RIKEN Natural Product Depository^47^, and among its top 10 most sensitive interactions with the diagnostic pool mutants were the *FLC2*, *DFG5*, *GAS1* and *HOC1^28^* genes. The *FCL2* product is a putative calcium channel involved in cell wall maintenance^48^, *DFG5* encodes a glycosylphosphatidylinositol (GPI)-anchored membrane protein required for cell wall biogenesis in bud formation^49^, *GAS1* encodes a β-1,3-glucanosyltransferase required for cell wall assembly^50–52^, and *HOC1* codes for an alpha-1,6-mannosyltransferase involved in cell wall mannan biosynthesis^53^. By comparing NP329’s diagnostic pool gene sensitivity profiles with the compendium of genetic interactions mapped in yeast and analyzing our data using the CG-TARGET software for chemical-genetic profile interpretation^28,54^, the top three high-confidence GO bioprocesses predicted to be perturbed by NP329 were “cell wall biogenesis” (GO:0042546), “cell wall organization or biogenesis” (GO:0071554), and “fungal-type cell wall organization or biogenesis” (GO:0071852). This strongly implicates the pseudojervine NP329 as a disrupter of proper cell wall biogenesis in yeast.

To further study this compound-process interaction, we hierarchically clustered the BIONIC PEG features, and we focused on the essential genes present in the **Fig. 2b** “glycosylation, protein folding/targeting, cell wall biosynthesis” bioprocess (**Fig. 6d**). We observed that 6 out of 16 NP329 sensitive essential genes lie in the bioprocess, as do 18 out of 20 BIONIC predicted sensitive essential genes. Within this bioprocess, BIONIC successfully predicts 4 (*BIG1*, *KRE5*, *KRE9*, *ROT1*) out of the 6 NP329 sensitive essential genes. These results indicate that BIONIC is able to both predict a relevant biological process targeted by the compound, and the specific sensitive genes. Moreover, the four sensitive genes successfully predicted by BIONIC were all closely clustered together based on the integrated BIONIC features (**Fig. 6**). *ROT1* encodes an essential chaperone required for N- and O-glycosylation in yeast^55^, and is required for normal levels of β-1,6-glucan^56^. Both *KRE5* and *BIG1* are also required for proper β-1,6 glucan synthesis^57,58^. These interactions further implicate that NP329 can interfere in the proper synthesis of β-1,6-glucan, an essential cell wall component. Since the chemical structure of NP329 is extremely similar to the natural product jervine, we tested the effect of jervine on the production of β-1,6-glucan. *KRE6* is a nonessential gene that, like its paralog *SKN1*, encodes a glucosyl hydrolase required for β-1,6-glucan biosynthesis^59^. We found that treatment of cells with 5 ug/mL of jervine reduced β-1,6-glucan levels to the same extent as a kre6 deletion mutant, likely by inhibiting *KRE6* and its paralog *SKN1* (**Fig. S6**). These results show that BIONIC can predict relevant chemical-genetic interactions, and has the potential to link compounds to their cellular targets.

## Discussion

We present BIONIC, a new deep-learning algorithm that extends the graph convolutional network architecture to integrate biological networks. BIONIC produces gene features that capture functional information well when compared to other unsupervised methods^12,17^ as determined by a range of benchmarks and evaluation criteria, covering a diverse set of downstream applications, such as gene co-annotation prediction, functional module detection, and gene function prediction. BIONIC can utilize labelled data in a semi-supervised fashion when it is available, and it can be purely unsupervised otherwise. BIONIC performs well for a range of numbers of input networks and network sizes, where established methods are not able to scale past relatively few networks or scale only with reduced performance.

From an application perspective, we demonstrated that BIONIC can be used to predict pairs of related genes (co-annotation prediction), identify functional gene modules (module detection), and accurately predict functional gene labels (gene function prediction). One of the main goals of this work is to generate fully integrated features encoding functional information for a particular organism, such that the resulting features can be used to predict numerous different aspects of cell and organism function^60^. As a proof-of-concept, we integrated 3 different yeast networks, incorporating protein-protein interaction, co-expression, and genetic interaction data. We also used BIONIC features to generate predictions for essential gene chemical sensitivities. We experimentally validated these chemical-genetic predictions, of which a significant number were correct - indicating BIONIC is effective at prediction and hypothesis generation. BIONIC is a general network integration algorithm, so in practice, input networks do not necessarily need to consist of genes or proteins. A potential application of BIONIC is patient network integration, where nodes represent patients and edges connect patients, weighted by the similarity of their profiles. These networks are often multi-modal, linking patients based on (for example) clinical, genomic or metabolomic information, thereby producing multiple, distinct networks^61,62^. BIONIC features representing the integration of these networks could, for instance, be clustered to identify groups of related patients that may have a particular disease subtype. Such subtypes may reflect distinct dysregulated cellular programs, potentially informing precision medicine treatments.

In a global sense, BIONIC performs well and captures relevant functional information across input networks. However, input networks do not have uniform quality and some networks may only describe certain types of functional relationships effectively (such as those within a particular biological process) while obscuring other relationships. Indeed, while BIONIC is able to capture a greater number of functional modules than a given input network alone (**Fig. 2c, Fig. S4**), BIONIC does not capture every functional module present in the input networks (**Fig. 2c, Fig. S4, Supplementary Data Files 4, 5**). This is likely due to some networks obscuring signals present in other networks. Implementing more advanced input network feature weighting should ensure that high-quality information is preferentially encoded in the learned features and that low-quality information is not. This may help to identify which functional relationships are driven by which networks and network types, thereby indicating which parts of the functional spectrum have good or poor coverage and identifying areas to target for future experimental work. Additionally, while BIONIC features are useful for identifying groups of related genes at a local level (complexes, pathways and bioprocesses), more assessment will be required to determine the utility of these features for capturing global characteristics of the input networks (such as centrality), and corresponding prediction tasks (such as gene essentiality prediction)^63^.

Interestingly, the naive union of networks approach performs surprisingly well, motivating its inclusion as a baseline in any network integration algorithm assessments. While the union network contains all possible relationships across networks, it likely contains relatively more false-positive relationships in the integrated result, since all false-positives in the input networks are retained by the union operation. Thus, the union should work well for high quality networks, but perform poorly with noisy networks.

Our chemical-genetic analysis demonstrates the potential of BIONIC to provide high resolution target predictions from limited experimental data. While BIONIC performs well at predicting essential gene chemical-genetic interactions, further improvements in performance could potentially be made through an optimized choice of input networks that specifically indicate these chemical sensitivities. Identifying or constructing such networks is a potential area of future work. BIONIC chemical-genetic interaction predictions could also be used to instead generate a set of putative non-sensitive genes for a given compound, indicating bioprocesses where the compound is not active. This would reduce the size of the experimental space when screening, resulting in more rapid and less expensive data generation. Finally, strong BIONIC chemical-genetic interaction predictions that are not reflected in the experimental data could indicate potential experimental false negatives that require additional investigation.

BIONIC learns gene features based solely on their topological role in the given networks. A powerful future addition to BIONIC would be to include gene or protein features such as amino acid sequence^64^, protein localization^65^, morphological defect^66^, or other non-network features to provide additional context for genes in addition to their topological role. Continued development of integrative gene function prediction using deep learning-based GCN and encoder-decoder technologies will enable us to map gene function more richly and at larger scales than previously possible.

## Acknowledgements

We thank B. Andrews, M. Costanzo and C. Myers for their insightful comments. We also thank M. Fey for adding important features to the Pytorch Geometric library for us. This work was supported by NRNB (U.S. National Institutes of Health, National Center for Research Resources grant number P41 GM103504). Funding for continued development and maintenance of Cytoscape is provided by the U.S. National Human Genome Research Institute (NHGRI) under award number HG009979. This work was also supported by CIHR (Canadian Institutes of Health Research) Foundation grant number FDN-143264, US National Institutes of Health grant number R01HG005853 and joint funding by Genome Canada (OGI-163) and the Ministry of Economic Development, Job Creation and Trade, under the program Bioinformatics and Computational Biology (BCB). This work was supported by the National Research Council of Canada through the AI for Design program. This work was supported by CIFAR AI Chair programs. This work was also supported by JSPS KAKENHI Grant numbers JP15H04483 (CB and YO), JP17H06411 (CB and YY), JP18K14351 (KIN), JP19H03205 (YO), JP20K07487 (DY), and a RIKEN Foreign Postdoctoral Fellowship (SCL).

## Author Contributions

DTF conceived of and developed the method and computational experiments. SCL and MY performed the chemical-genetic screens. ZL provided resources for the TS mutant collection. LAVI preprocessed and provided the chemical genetic data. HO provided the chemical matter and information about the screened compounds. SCL and ZL constructed the drug hypersensitive TS mutant collection. KIN, DN, and YO performed the jervine biochemical validation. DTF, SCL, YY, YO, BW, GDB, and CB wrote the manuscript. BW, GDB, and CB conceived of and supervised the project.

## Online Methods

### BIONIC Method Overview

An undirected input network can be represented by its adjacency matrix *A* where *A_ij_* = *A_ji_* > 0 if node *i* and node *j* share an edge and *A_ij_* = *A_ji_* = 0otherwise. BIONIC first preprocesses each input network to contain the union of nodes across all input networks and ensures the corresponding row and column orderings are the same. In instances where networks are extended to include additional nodes not originally present in them (so all input networks share the same union set of nodes), the rows and columns corresponding to these nodes are set to 0.

BIONIC encodes each input network using instances of a GCN variant known as the Graph Attention Network (GAT)^27^. We selected this architecture because of its considerable performance improvements over existing architectures on a range of node classification tasks^27^. The GAT has the ability to learn alternative network edge weights, allowing it to downweight or upweight edges based on their importance for the network reconstruction task. In the original formulation, the GAT assumes binary network inputs. We modify the GAT to consider *a priori* network edge weights. The GAT formulation is then given by:

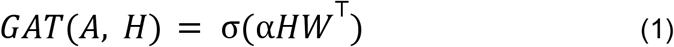

where

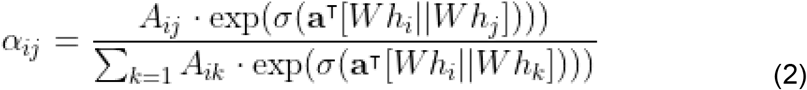

Here, *W* is a trainable weight matrix which projects aggregated node features into another feature space, **a** is a vector of trainable attention coefficients which determine the resulting edge weighting, *h_i_* is the feature vector for node *i* (that is, the *i*th row of feature matrix *H*), ||denotes the concatenation operation and σ corresponds to a nonlinear function (in our case a leaky rectified linear unit (LeakyReLU)) which produces more sophisticated features than linear maps. (1) corresponds to a node neighborhood aggregation and projection step which incorporates an edge weighting scheme (2). In practice, several edge weighting schemes (known as attention heads) are learned and combined simultaneously, resulting in:

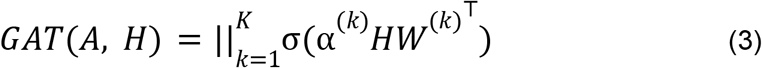

where *K* is the number of attention heads. This is done to stabilize the attention learning process, as per the author’s original results^27^. In our experiments we use 10 attention heads per GAT encoder, each with a hidden dimension of 68, as per our hyperparameter optimization results (see **Obtaining Integrated Results**, **Supplementary Data File 1**).

Initial node features *H_init_* are one-hot encoded so that each node is uniquely identified (i.e. *H_init_* = *I* where *I* is the identity matrix). These features are first mapped to a lower dimensional space through a learned linear transformation to reduce memory footprint and improve training time. BIONIC encodes each network by passing it through several sequential GAT layers to learn node features based on higher-order neighborhoods. Outputs from each GAT pass are then summed to produce the final network-specific features (**Fig. S1**). Based on the hyperparameter optimization results, we used three GAT layers in our experiments. After all networks are separately encoded, the network-specific node features are combined through a weighted, stochastically masked summation given by:

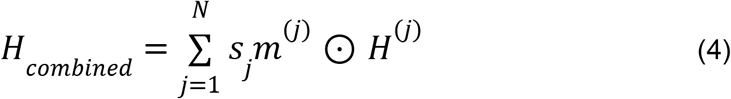

Here, *N* is the number of input networks, *s_j_* is the learned scaling coefficient for feature representations of network *j*, ⊙ is the element-wise product, *H*^(*j*)^ is the matrix of learned feature vectors for nodes in network *j*, and *m*^(*j*)^ is the node-wise stochastic mask for network *j*, calculated as:

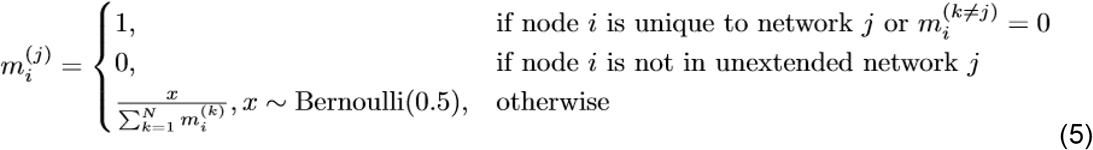

The mask *m* is designed to randomly drop node feature vectors produced from networks with the constraint that a node cannot be masked from every network, and node features from nodes not present in the original, unextended networks are dropped. This masking procedure forces the network encoders to compensate for missing node features in other networks, ensuring the encoders learn cross-network dependencies and map their respective node features to the same feature space. The network scaling vector *s* in (5) enables BIONIC to scale features in a network-wise fashion, affording more flexibility in learning the optimal network-specific node features for the combination step. *s* is learned with the constraint that its elements are positive and sum to 1, ensuring BIONIC does not over- or negatively-scale the features.

To obtain the final, integrated node features *F*, BIONIC maps *H_combined_* to a low dimensional space through a learned linear transformation. In *F*, each column corresponds to a specific learned feature and each row corresponds to a node. To obtain a high quality *F*, BIONIC uses an unsupervised training objective. When gene labels are provided, an additional semi-supervised training objective is also used.

For the unsupervised training objective, BIONIC decodes *F* into reconstructions of the original input networks and minimizes the discrepancy between the reconstructions and the inputs. The decoded network reconstruction is given by:

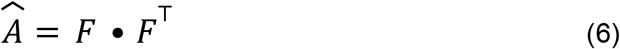

The unsupervised loss is then given by:

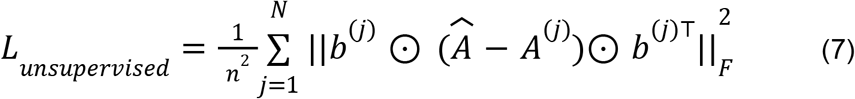

where *n* is the total number of nodes present in the union of networks, *b*^(*j*)^ is a binary mask vector for network *j* indicating which nodes are present (value of 1) or extended (value of 0) in the network, *A*^(*j*)^ is the adjacency matrix for network *j* and ∥ · ∥_*F*_ is the Frobenius norm. This loss represents computing the mean squared error between the reconstructed network 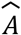 and input *A*^(*j*)^ while the mask vectors remove the penalty for reconstructing nodes that are not in the original network *j* (i.e. extended), then summing the error for all networks.

For the semi-supervised training objective, BIONIC first predicts gene labels by mapping *F*to a matrix of class predictions as follows:

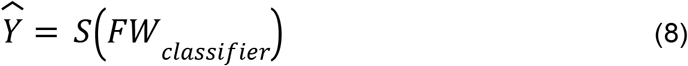

where *S*is the sigmoid function and *W_classifier_* is a trainable weight matrix. The resulting class prediction matrix 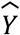 has genes as rows and class labels as columns. The ground-truth label matrix *Y* indicates the correct labels for a set of genes in the input networks. *Y*is extended to include zero vectors for any genes present in the input networks but not present in the labels, ensuring it has the same shape as 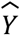. The semi-supervised loss is then given by:

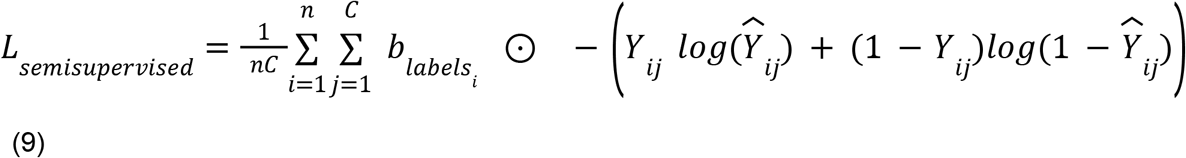

where *n* is the total number of nodes present in the union of networks, *C*is the number of classes, 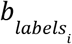 is a binary mask indicating whether node *i* was present in the original label set (value of 1) or was extended (value of 0). *log*indicates the natural logarithm. This loss represents the masked binary cross entropy between the predicted labels 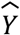 and the true labels *Y* ignoring the loss of any nodes not originally present in *Y*.

The final loss BIONIC trains to minimize is a weighted sum of the unsupervised and semi-supervised losses:

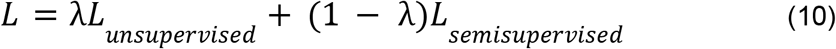

where λ is a value in the range [0, 1] indicating the relative weights of the two losses. When no labelled data is available, λ is set to 1.

### Implementation Details

BIONIC was implemented using PyTorch^67^, a popular Python-based deep learning framework, and relies on functions and classes from the PyTorch Geometric library^68^. It uses the Adam^69^ optimizer to train and update its weights. To be scalable in the number of networks, BIONIC utilizes an optional network batching approach where subsets of networks are sampled and integrated at each training step. The sampling procedure is designed so that each network is integrated exactly once per training step. Network batching yields a constant memory footprint at the expense of increased runtime with no empirical degradation of feature quality. This feature is provided for additional scalability over what is demonstrated in this work, and was not used in any of our reported experiments. Additionally, BIONIC is scalable in the number of network nodes. It uses a node sampling approach (equivalent to mini-batch training, where nodes are samples) to learn features for subsets of nodes in a network, and a neighborhood sampling procedure to subsample node neighborhoods. Node sampling ensures only part of a network needs to be retained in memory at a time while neighborhood sampling reduces the effective higher order neighborhood size in sequential GAT passes, again reducing the number of nodes required to be retained in memory at any given time - further reducing BIONIC’s memory footprint.

For very large networks where the initial node feature matrix (i.e. the identity matrix) cannot fit into memory due to limitations with PyTorch matrix operations, BIONIC incorporates a singular value decomposition (SVD) based approximation. First, the union of networks is computed by creating a network that contains the nodes and edges of all input networks. If an edge occurs in multiple networks, the maximum weight is used. A low-dimensional SVD approximation of the normalized Laplacian matrix of the union network is computed and used as the initial node features for each network. Finally, BIONIC uses sparse representations of network adjacency matrices (except for the input node feature matrix, see above), further reducing memory footprint. All BIONIC experiments in this paper were run on an NVIDIA Titan X GPU with 12GB of VRAM, no more than 16GB of system RAM and a single CPU.

### Network Preprocessing

The yeast protein-protein interaction network^5^ and human protein-protein interaction networks^3,6,7,41^ were obtained from BioGRID^70^, genetic interaction profiles^11^ were obtained directly from the published supplementary data of Costanzo et al. 2016, and gene expression profiles^9^ were obtained from the SPELL database^8^. These networks were chosen since they had the most functional information compared to other networks in their class (i.e. protein-protein interaction networks, co-expression networks, and genetic interaction networks). To create a network from the genetic interaction profiles, genes with multiple alleles were collapsed into a single profile by taking the maximum profile values across allele profiles. Pairwise Pearson correlation between the profiles was then calculated, and gene pairs with a correlation magnitude greater than or equal to 0.2 were retained as edges, as established^11^. For the gene expression profiles, networks were constructed by retaining gene pairs with a profile Pearson correlation magnitude in the 99.5th percentile. Co-expression and genetic interaction networks had their edge weights normalized to the range [0, 1].

### Obtaining Integrated Results

The naive union of networks benchmark was created by taking the union of node sets and edge sets across input networks. For edges common to more than one network, the maximum weight was used. For all other methods, automated hyperparameter optimization was performed to ensure hyperparameters were chosen consistently and fairly. Here, a Schizosaccharomyces pombe genetic interaction network^34^, co-expression network^33^, and protein-protein interaction network^32^ were used as inputs to the integration methods. To perform one iteration of the hyperparameter optimization, a random combination of hyperparameters was uniformly sampled over a range of reasonable values for each method and used to integrate the three pombe networks. The integration results were then evaluated using a pombe protein complex standard (obtained from https://www.pombase.org/data/annotations/Gene_ontology/GO_complexes/Complex_annotation.tsv). The evaluations consisted of a co-annotation prediction, module detection, and gene function prediction assessment (see **Evaluation Methods**). This procedure was repeated for 50 combinations of hyperparameters, for each method. For methods that produced features (deepNF, Mashup, multi-node2vec and BIONIC), a feature dimension of 512 was used to ensure results were comparable across methods. For methods which required a batch size parameter (deepNF and BIONIC), the batch size was set to 2048 to ensure reasonable computation times. Hyperparameter combinations were then ranked for each method across the three evaluation types and the hyperparameter combination corresponding to the highest average rank across evaluation types was chosen. The hyperparameter optimization results are found in **Supplementary Data File 1**. Note that the Union method was not included in the hyperparameter optimization because it has no hyperparameters. Additionally, the Mashup method used 44 hyperparameter combinations rather than 50, since 6 hyperparameter combinations exhausted the available memory resources and did not complete.

All integration results reported were obtained by integrating networks using the set of hyperparameters identified in the hyperparameter optimization procedure. BIONIC features used in the **Fig. 2, 3, 6** analyses are found in **Supplementary Data File 8**. Co-annotation prediction, module detection, and gene function prediction standards used in **Fig. 2**-**5** are found in **Supplementary Data File 9**.

### Benchmark Construction

Functional benchmarks were derived from GO Biological Process ontology annotations, KEGG pathways and IntAct complexes for yeast, and CORUM complexes for human (**Supplementary Data File 3**). Analyses were performed using positive and negative gene pairs, clusters or functional labels obtained from the standards as follows: the GO Biological Process benchmark was produced by filtering IEA annotations, as they are known to be lower quality, removing genes with dubious open reading frames, and filtering terms with more than 30 annotations (to prevent large terms, such as those related to ribosome biogenesis, from dominating the analysis^71^). For the co-annotation benchmark, all gene pairs sharing at least one annotation were retained as positive pairs, while all gene pairs not sharing an annotation were considered to be negative pairs. KEGG, IntAct and CORUM benchmarks were produced analogously, without size filtering.

For the module detection benchmark, clusters were defined as the set of genes annotated to a particular term, for each standard. Modules of size 1 (singletons) were removed from the resulting module sets as they are uninformative. For the per-module analyses in **Fig. 2c**, **S3, S4**, and **Supplementary Data File 4, 5** we also removed any modules of size 2 since these modules had highly variable Jaccard scores.

The supervised standards were obtained by treating each gene annotation as a class label, leading to genes with multiple functional classes (i.e. a multilabel classification problem). The standards were filtered to only include classes with 20 or more members for GO Biological Process and KEGG, or 10 members for IntAct. This was done to remove classes with very few data points, ensuring more robust evaluations.

The granular function standard in **Fig. 2b, 3b, 6** was obtained from the Costanzo et al. 2016 supplementary materials. Any functional category with fewer than 20 gene members was removed from the analysis to ensure only categories with robust evaluations were reported.

### Evaluation Methods

We used a precision-recall (PR) based co-annotation framework to evaluate individual networks and integrated results. We used PR instead of receiving operator curve (ROC) because of the substantial imbalance of positives and negatives in the pairwise benchmarks for which ROC would overestimate performance. Here, we computed the pairwise cosine similarities between gene profiles in each network or integration result. Due to the high-dimensionality of the datasets, cosine similarity is a more appropriate measure than Euclidean distance since the contrast between data points is reduced in high-dimensional spaces under Euclidean distance^72^. PR operator points were computed by varying a similarity threshold, above which gene or protein pairs are considered positives and below which pairs are considered negative. Each set of positive and negative pairs was compared to the given benchmark to compute precision and recall values. To summarize the PR curve into a single metric, we computed average precision (AP) given by:

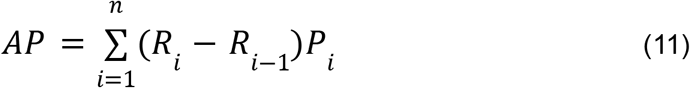

where *n* is the number of operator points (i.e. similarity thresholds) and *P_i_* and *R_i_* are the precision and recall values at operator point *i* respectively. This gives the average of precision values weighted by their corresponding improvements in recall. We chose this measure over the closely related area under the PR curve (AUPRC) measure since AUPRC interpolates between operator points and tends to overestimate actual performance^73^.

The module detection evaluation was performed by clustering the integrated results from each method and comparing the coherency of resulting clusters with the module-based benchmarks. Since the benchmarks contain overlapping modules (i.e. one gene can be present in more than one module) which prevents the use of many common clustering evaluation metrics (since these metrics assume unique assignment of gene to cluster), the module sets are subsampled during the evaluation to ensure there are no overlapping modules (the original module sets are used as-is for the per-module-optimized experiments in **Fig. S4, Supplementary Data File 5**). Next, the integrated results are hierarchically clustered with a range of distance metrics (Euclidean and cosine), linkage methods (single, average and complete) and thresholds to optimize benchmark comparisons over these clustering parameters (this is done for all methods that are compared). The resulting benchmark-optimized cluster sets are compared to the benchmark module sets by computing adjusted mutual information (AMI) - an information theoretic comparison measure which is adjusted to normalize against the expected score from random clustering. The highest AMI score for each integration approach is reported - ensuring the optimal cluster set for each dataset across clustering parameters is used for the comparison and that our results are not dependent on clustering parameters. Finally, this procedure is repeated ten times to control for differences in scores due to the cluster sampling procedure. The sets of clustering parameter-optimized BIONIC clusters obtained from the **Fig. 2** integration for each standard are in **Supplementary Data File 4**.

To perform the supervised gene function prediction evaluation, ten trials of five-fold cross validation were performed using support vector machine (SVM) classifiers each using a radial basis function kernel^31^. The classifiers were trained on a set of gene features obtained from the given integration method with corresponding labels given by the IntAct, KEGG and GO Biological Process supervised benchmarks in a one-versus-all fashion (since each individual gene has multiple labels). Each classifier’s regularization and gamma parameters were tuned in the validation step. For each trial, the classifier results were evaluated on a randomized held out set consisting of 10% of the gene features not seen during training or validation and the resulting classification accuracy was reported.

The granular functional evaluations in **Fig. 2b, 3b** were generated by computing the average precision (as mentioned in the precision-recall evaluation framework description) for the gene subsets annotated to the given functional categories.

To perform the module comparison analysis in **Fig. 2c**, we additionally applied the module detection analysis performed in **Fig. 2a** to the input networks. Here, the interaction profiles of the networks were treated as gene features and the clustering parameters were optimized to best match the IntAct complexes standard. We compared the resulting module sets from the input networks and BIONIC features to known protein complexes given by the IntAct standard. For each complex in the standard, we reported the best matching predicted module in each dataset as determined by the overlap (Jaccard) score between the module and the known complex (**Supplementary Data File 4**). To generate the Venn diagram, we defined a complex to have been captured in the dataset if it had an overlap score of 0.5 or greater with a predicted module.

To perform the LSM2-7 module analysis in **Fig. 2d**, we analyzed the predicted module in each dataset that had the highest overlap score with the LSM2-7 complex. We created a network from the BIONIC features by computing the cosine similarity between all pairs of genes and setting all similarities below 0.5 to zero. The resulting non-zero values were then treated as weighted edges to form a network. We extracted a subnetwork from each of the protein-protein interaction, co-expression, genetic interaction and newly created BIONIC networks, consisting of the best scoring predicted module and the genes showing direct interactions with those in the predicted module. We laid out these networks using the spring-embedded layout algorithm in Cytoscape^74^. The edges in the protein-protein interaction network correspond to direct, physical interactions, and the edges in the co-expression and genetic interaction networks correspond to the pairwise Pearson correlation of the gene profiles, as described above.

To perform the semi-supervised network integration experiment in **Fig. 4**, we first generated randomized train and test sets. Here, 20% of genes were randomly held out in each gene function benchmark (IntAct, KEGG, and GO Biological Process) separately, and retained for downstream evaluations. These benchmarks consist of functional labels for a set of yeast genes (protein complex membership in IntAct, pathway membership in KEGG, and biological process annotation in GO Biological Process), and are the same benchmarks used in the gene function prediction evaluation (**Fig. 2a, 3a**). The remaining 80% of genes were used for training GeneMANIA and BIONIC. To generate test sets for the co-annotation prediction benchmarks, we removed any co-annotations where both genes were present in the training set. To generate test sets for the module detection benchmarks, we removed any modules consisting entirely of genes in the training set. We then integrated the three yeast networks from the **Fig. 2, 3** analysis (a protein-protein interaction^5^, gene co-expression^9^, and genetic interaction network^11^) using the supervised GeneMANIA, BIONIC without using any labelled data (unsupervised), and a semi-supervised mode of BIONIC which uses the labelled data (semi-supervised). Each integration result was then evaluated using the held-out test data. For the co-annotation prediction and module detection evaluations, the integrated features from BIONIC (both unsupervised and semi-supervised), and the integrated network from GeneMANIA were evaluated. Both GeneMANIA and the semi-supervised BIONIC generate gene label predictions directly, without the need for an additional classifier like in the **Fig. 2a** gene function prediction evaluation. However, the unsupervised BIONIC does not generate gene label predictions (since it is given no labelled information to begin with). To ensure a consistent comparison with GeneMANIA and the semi-supervised BIONIC, we trained a classification head on top of the unsupervised BIONIC. The classification head architecture is identical to the semi-supervised BIONIC classification head, however, in the unsupervised case we only allow gradients from the classification loss objective to backpropagate to the classification head, not the rest of the model. This ensures a comparable classification model can be trained on top of the unsupervised BIONIC model, without the labelled data affecting the model weights like in the semi-supervised case. GeneMANIA does not generate multi-label predictions, and so we used GeneMANIA to generate label predictions for each class individually and then performed Platt scaling to convert these binary class predictions to multi-label predictions^17,75^. The gene function prediction evaluations were then performed by comparing the gene label predictions from the integration methods, to the held-out test labels. This entire procedure, starting with the train-test set partitioning, to the final evaluations, was repeated a total of 10 times to control for performance variability due to the partitioning procedure.

### Network Scaling Experiment

To perform the network scaling experiment, we uniformly sampled subsets of the yeast co-expression networks (**Supplementary Data File 2**). We performed 10 integration trials for each network quantity, and these trials were paired (i.e. each method integrated the same randomly sampled sets of networks). The average precision scores of the resulting integrations with respect to the KEGG pathways co-annotation standard (**Supplementary Data Files 3**) were then reported. The Mashup method did not scale to the 7 network input size or beyond on a machine with 64GB of RAM.

### Node Scaling Experiment

The node scaling experiment was performed by uniformly subsampling the nodes of four large human protein-protein interaction networks^3,6,7,41^ (**Supplementary Data File 2**) for a range of node quantities and integrating these subsampled networks. Ten trials of subsampling were performed for each number of nodes (paired, as above) and the average precision scores with respect to the CORUM complexes co-annotation standard (**Supplementary Data File 3**) were reported. The Mashup method did not scale to 4000 nodes or beyond on a machine with 64GB of RAM.

### Gene-Chemical Sensitivity Predictions

Chemical-genetic profiles against a diagnostic set of 310 non-essential yeast gene deletion mutants were obtained from a previous study^28^. The genes were chosen using the COMPRESS-GI algorithm, which selected a set of 157 genes capturing a majority of the functional information within genome-wide genetic interaction data^76^, along with 153 genes that were manually selected to complement the set. Haploid deletion mutants for the gene set were constructed in a genetic background that conferred drug hypersensitivity (*pdr1Δpdr3Δsnq2Δ*) using synthetic genetic analysis (SGA) technology, and each mutant strain was barcoded with a unique 20 bp DNA identifier adjacent to a common priming site. The mutant collection was grown and stored as a pooled library in YPD-glycerol (15% v/v). A set of approximately 10,000 compounds from the RIKEN Natural Product Depository (NPDepo) were interrogated. Screens were done in 96-well format, where a single well contained the entire pool of 310 mutants at a density of 4.65×10^5^ cells/mL and 196 uL of YPGal media (1% yeast extract, 2% peptone, 2% galactose). Each well was treated with 2 uL of compound (1 mg/mL stock dissolved in DMSO). After 48 hours of growth in 30 C, genomic DNA was extracted from each compound-treated pool with an automated high-throughput nucleic acid purification robot (QIAcube HT, Qiagen). Mutant-specific barcodes and well-specific index tags were PCR-amplified using multiplex primers and a communal U2 primer. PCR products were pooled in 768-plex and gel-purified from 2% agarose gels using a Geneclean III kit. Amplicons were quantified using a Kapa qPCR kit, and were sequenced with an Illumina Hiseq 2500 machine at the RIKEN Center for Life Science Technologies. Sequencing data was processed using the BEAN-counter software^46^, which generated chemical-genetic interaction z-scores normalized against DMSO-only (1% DMSO) treated samples. False discovery rates (FDR) were estimated for biological process^11^ predictions, for each compound, and those compounds with an FDR < 25% were retained, resulting in a set of 1522 compounds and 289 genes (high confidence set)^28^. Next, interquartile range (IQR) scores were calculated from the chemical-genetic scores as follows:

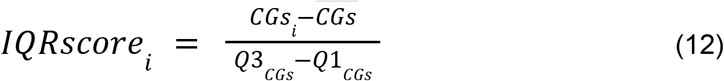

Here, *CGs_i_* is the chemical-genetic score for the *i*th replicate, 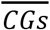 is the median of all chemical-genetic scores, *Q*3_*CGs*_ is the 75th percentile of chemical-genetic scores, and *Q*1_*CGs*_ is the 25th percentile of chemical-genetic scores. Tukey’s test^77^ was used to determine outliers based on the interquartile range of the distribution of IQR scores in the screen. Genes with at least one replicate that had a negative (sensitive) chemical-genetic score more than three times the interquartile range of the compound profile (i.e. “outlier” genes) were retained.

To predict chemical-genetic interactions using BIONIC, we first selected a set of 50 compounds to generate predictions on and experimentally validate. For each diagnostic pool compound, we filtered out any genes not present in the integrated BIONIC features (the same features used for the **Fig. 2, 3** analyses, referred to as PEG features). Any compounds with fewer than 2 outlier sensitive genes were then removed. For each of the remaining compounds, we randomly split the sensitive genes into train and test sets. Next, for a given compound, we computed BIONIC predictions for the test set genes. We did this by averaging the corresponding BIONIC PEG features for each gene in the training set under a cosine distance metric to get a representative feature vector in gene feature space for the given compound. The BIONIC predictions for the compound were then obtained by identifying the top 20 nearest genes to this feature vector (excluding genes in the training set).

To obtain a score for the BIONIC predictions, an ordered Fisher’s exact test was performed between the test set genes and the BIONIC predictions as follows:

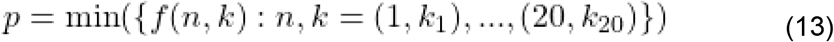

where

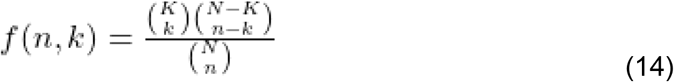

*p* corresponds to the minimum p-value obtained for progressively larger subsets of BIONIC’s 20 predictions, starting from the top prediction to the full set of 20 predictions. *n* is the number of total predictions made by BIONIC, and *k* is the number of those predictions that are correct. *k*_*i*_ corresponds to the number of correct predictions for the first *i* genes in the BIONIC predictions. *f* is the probability mass function of the hypergeometric distribution. Here, *K* corresponds to the number of genes found to be sensitive to the given compound. *N* is the total number of yeast essential genes in the analysis, specifically, essential genes for which TS mutants could be made and are also present in the BIONIC features (847 total genes). We chose the ordered Fisher’s exact test over the commonly used unordered version because BIONIC produces a ranked list of predictions. Taking into account the ordering of BIONIC predictions is a more fair assessment, since, for example, a compound may only have a small number of sensitive genes (fewer than 20). In this case, BIONIC’s top predictions may include these essential genes, however an unordered Fisher’s exact test would not consider this ranking and treat the full set of 20 predictions as equivalent, whereas the ordered test would consider the ranking.

The above process was repeated 5 times for new randomly sampled train and test gene splits, or up to the maximum number of train-test splits possible for compounds with fewer than 5 sensitive genes. Final p-values were obtained for each compound by averaging over the p-values from each trial. Compounds were ranked by most significant p-values and the top 50 compounds were selected for further screening. Sensitive essential gene predictions for a given compound were then generated by using the full set of sensitive diagnostic pool genes as the training set, computing a representative compound feature vector by averaging the training set BIONIC gene features, and identifying the top 20 nearest essential genes to this compound feature vector.

The BIONIC gene-chemical sensitivity predictions were benchmarked against experimental data obtained from chemical-genetic screens using a collection of temperature sensitive mutants for essential genes. We previously constructed a drug-hypersensitive, barcoded set of temperature sensitive (TS) mutants for 1181 TS alleles spanning 837 essential genes^45^. Similar to the diagnostic set of non-essential genes, this collection also contained the pdr1Δpdr3Δsnq2Δ triple deletion, and a 20 bp barcode was inserted next to a common priming site upstream of a natMX cassette integrated at the pdr3Δ locus. We conducted chemical-genetic screens against the 50 compounds initially selected for BIONIC analysis using the same method that was used to generate the diagnostic set profiles, except that the temperature sensitive mutant pools were incubated at 25 C instead of 30 C for 48 hours. We calculated chemical-genetic interaction Z-scores (CG scores) and removed non-specific technical effects using BEAN-counter software^46^. IQR scores were calculated as described above. Negative (sensitive) interactions that were more than four times the interquartile range (classified as “far outliers”) were used to validate the gene-chemical sensitivities predicted by BIONIC.

The significance of BIONIC sensitive essential gene predictions for each compound was determined by using an ordered Fisher’s exact test, as detailed above. The Benjamini-Hochberg procedure^78^ was applied to the resulting p-values at a false discovery rate of 5%.

To generate biological process^11^ predictions as reported in **Fig. 6a, b**, a Fisher’s exact test was performed between the full set of 20 BIONIC gene predictions and biological process gene annotations. We used the same annotations as in **Fig. 2b, 3b**^11^. If the BIONIC sensitive gene predictions were enriched for one or more bioprocesses, and these bioprocesses overlapped with the annotated bioprocess of the true sensitive genes, we considered this a correct bioprocess prediction. To generate the random benchmark in **Fig. 6b**, the gene labels of the BIONIC integrated features were randomly permuted and new essential sensitive gene predictions for the 50 selected compounds were generated in the same manner as the original BIONIC predictions (detailed above). This process was repeated for 1000 random gene label permutations to generate the benchmark distributions. The circle plot in **Fig. 6d** was produced by first hierarchically clustering the integrated BIONIC gene features, subsetted to essential genes annotated to the glycosylation, protein folding/targeting, cell wall biosynthesis bioprocess. Two clustering thresholds were chosen to generate clusters - broadly indicating the hierarchical organization of the BIONIC gene features. The first, most granular clustering threshold was adaptively chosen to generate clusters best matching known protein complexes, as defined by the IntAct Complexes standard^36^. For each protein complex in the standard, the clustering threshold was optimized to produce the cluster best matching this protein complex. For clusters not matching known complexes, the largest complex optimized threshold was used. The second, higher clustering threshold was set to a cophenetic distance of 0.9.

The BIONIC essential gene sensitivity predictions can be found in **Supplementary Data File 7**.

### Quantification of β-1,6-glucan levels

Wild-type (*his3Δ* in the BY4741 background) and the *kre6Δ* strain (YOC5627) of *S*. *cerevisiae* were grown in YPD at 25°C with shaking at 200 rpm to 1 × 10^7^ cells/mL. Wild-type cells were treated with 5 μg/mL jervine (J0009; Tokyo Chemical Industry, Tokyo, Japan) for 4 hrs. The samples were centrifuged at 15,000 × *g* for 3 minutes, and the supernatant was discarded. The pellet was washed and suspended in PBS, adjusted to 1 × 10^6^ cells/mL, and autoclaved for 20 min. After centrifugation at 15,000 × *g* for 1 minute, the supernatant was stored on ice (Sample A) and the pellet was further extracted. The β-1,6-glucan was extracted from the pellet using a slightly modified version of the protocol of Kitamura et al. (2009)^79^. First, 500 mL of 10% TCA was added to the culture, which was incubated on ice for 10 minutes. After centrifugation at 15,000 × *g* for 3 minutes, the samples were washed twice with DW. The pellet was suspended in 500 μL of 1 N NaOH and incubated at 75°C for 1 hour. The solution was mixed with 500 μL of 1 M HCl and Tris buffer (10 mM Tris-HCl, pH 7). After centrifugation at 15,000 × *g* for 1 minute, the supernatant was stored on ice (Sample B). The total amounts of β-1,6-glucan in Samples A and B were measured according to the method of Yamanaka et al. (2020)^80^.

### Data Availability

All data, standards and BIONIC yeast features are available in **Supplementary Data Files 7, 8**.

### Code Availability

The BIONIC code is available at https://github.com/bowang-lab/BIONIC.

## Supplementary Figures

**Figure S1.**
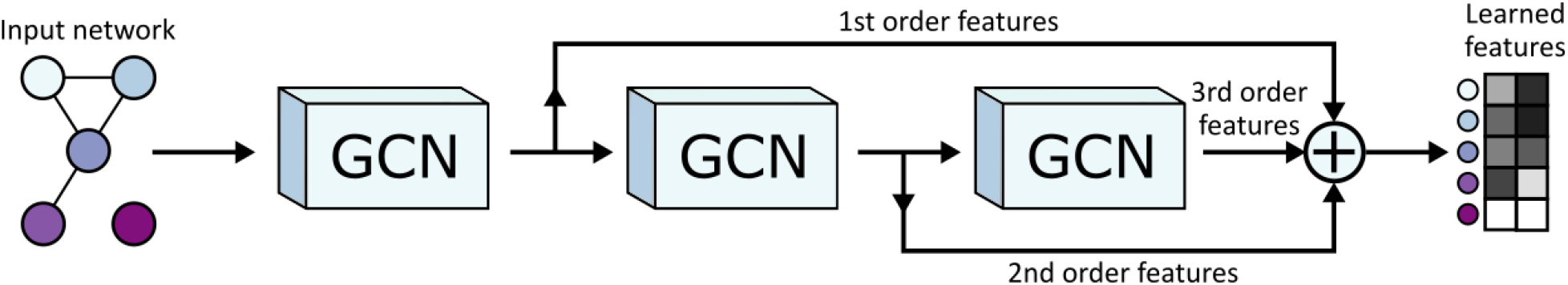
A more detailed view of an individual network encoder, including residual connections. A network specific graph convolutional network is used to encode the input network for increasing neighborhood sizes. The first GCN in the sequence learns features for a given node based on the node’s immediate neighborhood (1st order features). The next GCN learns features based on the node’s second order neighborhood (2nd order features), and so on. The node feature matrices learned by each GCN pass are summed together to create the final learned, network-specific features. Summing the outputs of the various GCNs in this way creates residual connections, allowing features from multiple neighborhood sizes to generate the final learned features, rather than just the final neighborhood size. This figure shows three GCN layers, but BIONIC uses the same pattern of connections for any number of GCN layers. Note that the GCN layers for a given encoder share their weights, so in effect, there is a single GCN layer for each encoder.

**Figure S2.**
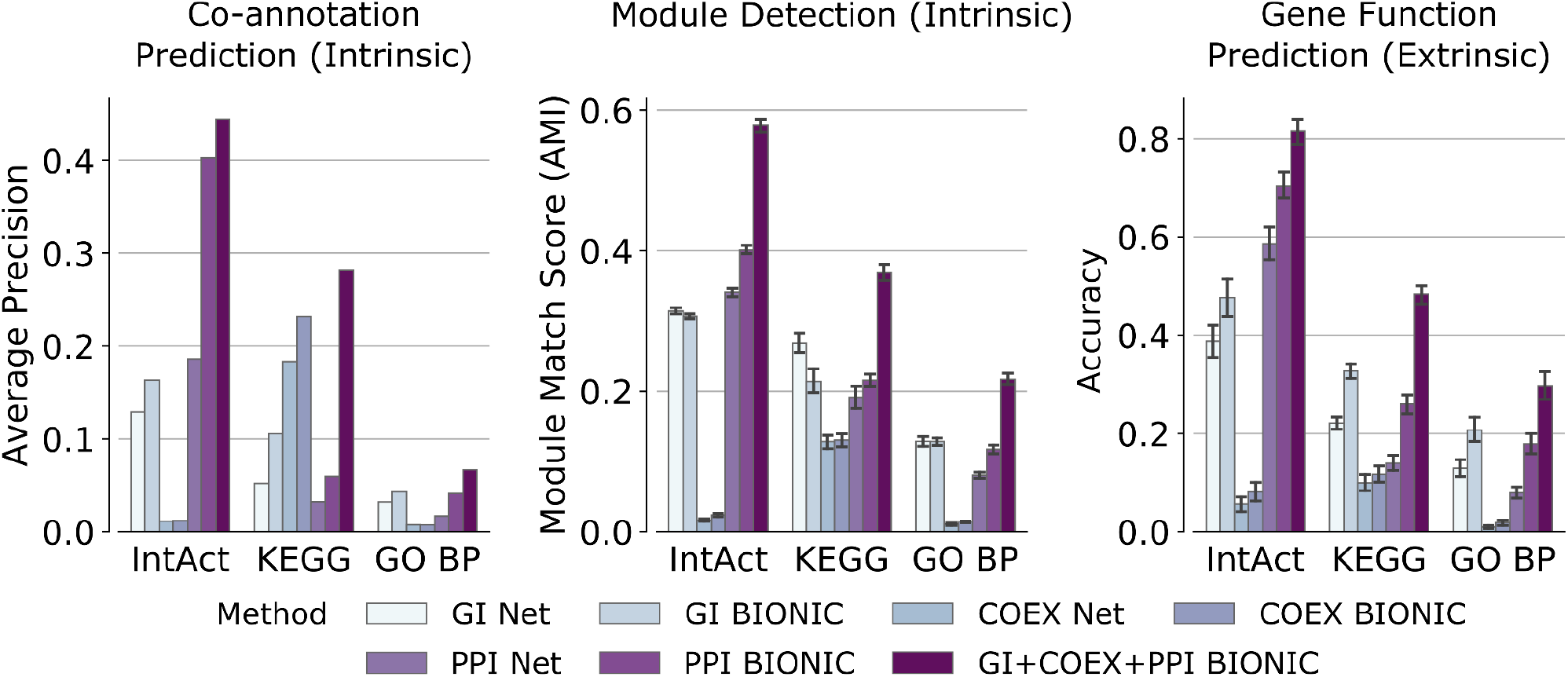
A comparison of individual networks (denoted “Net”), their corresponding features encoded using BIONIC (denoted “BIONIC”), as well as the BIONIC integration of these networks (denoted “GI+COEX+PPI BIONIC”). BP = Biological Processes, GI = Genetic Interaction, COEX = Co-expression, PPI = Protein-protein Interaction. These are the same networks and evaluations used in **Fig. 2**. Error bars indicate the 95% confidence interval.

**Figure S3.**
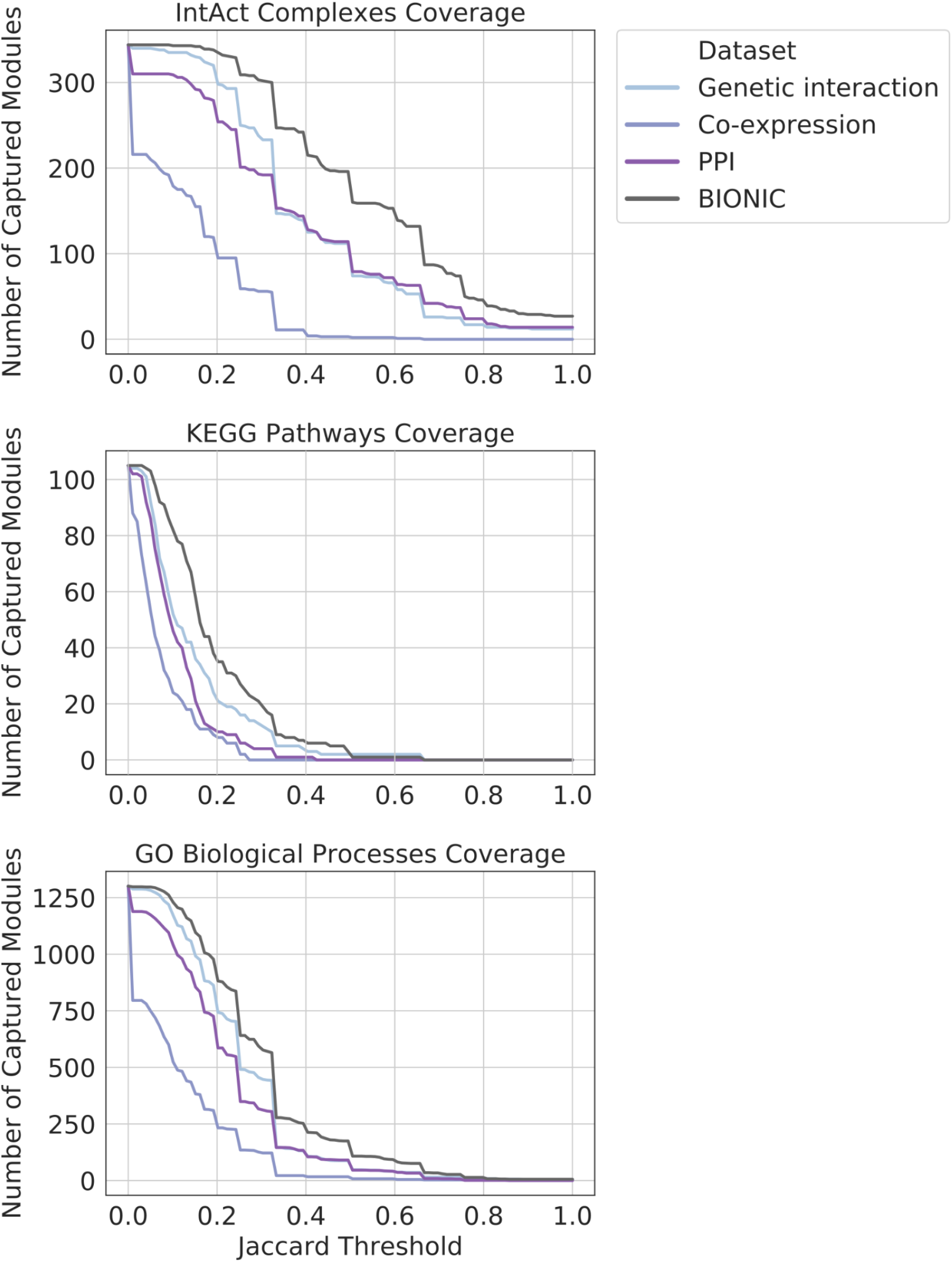
Coverage of functional gene modules by individual networks and the BIONIC integration of these networks (denoted BIONIC), as determined by a parameter optimized module detection analysis where the clustering parameters were optimized for each module individually. The number of captured modules is reported for a range of overlap scores (Jaccard threshold). Higher threshold indicates greater correspondence between the clusters obtained from the dataset and their respective modules given by the standard. PPI = protein-protein interaction. These are the same networks and BIONIC features as **Fig. 2**.

**Figure S4.**
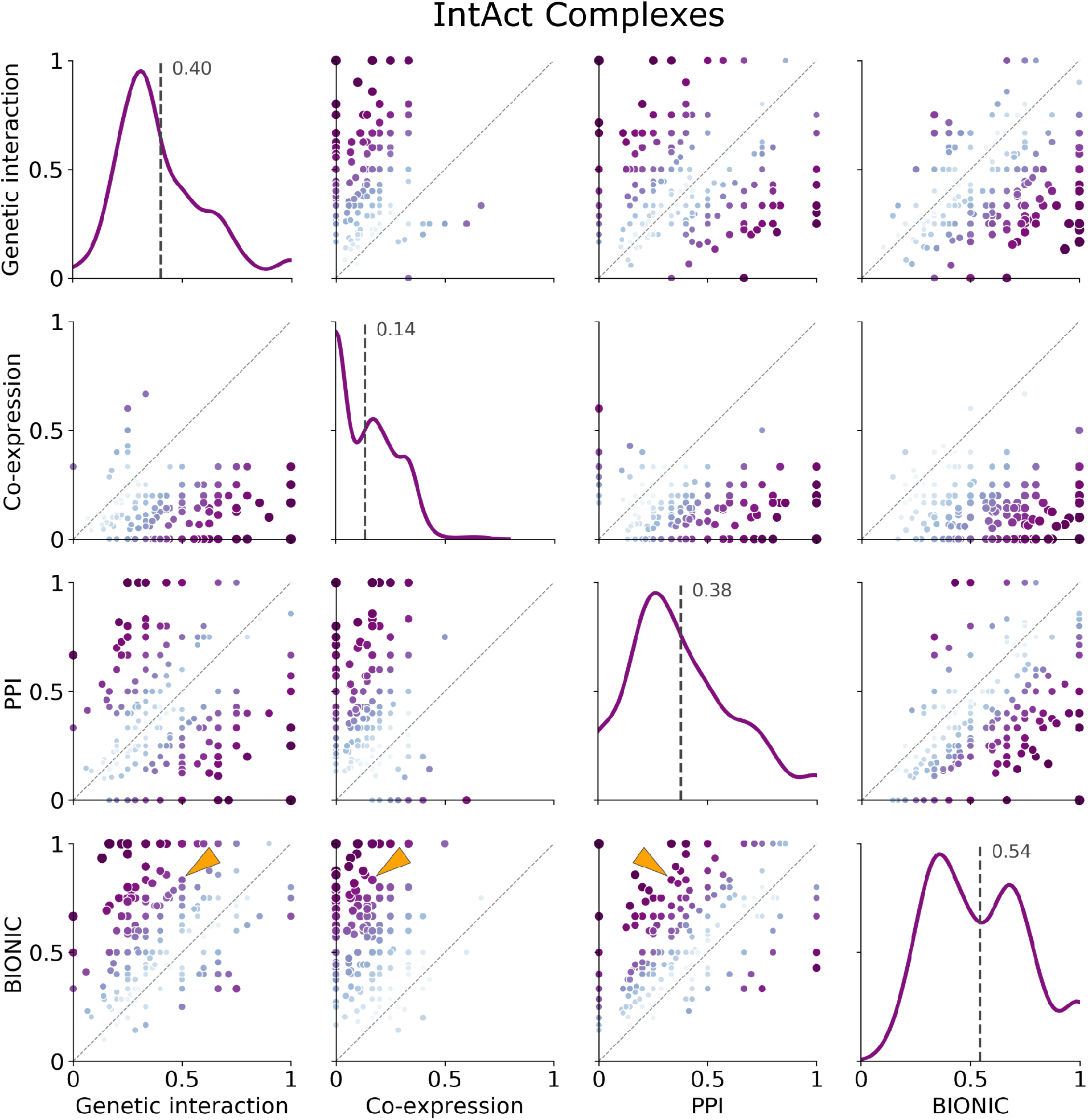
Known protein complexes (as defined by the IntAct standard) captured by individual networks and the BIONIC integration of these networks (denoted BIONIC). Hierarchical clustering was performed on the datasets and resulting clusters were compared to known IntAct complexes and scored for set overlap using the Jaccard score (ranging from 0 to 1). The clustering algorithm parameters were optimized for each module individually, unlike the analysis in **Fig. 2** where the clustering parameters were optimized for the standard as a whole. Each point is a protein complex, as in **Fig. 2c**. The dashed line indicates instances where the given data sets achieve the same score for a given module. Histograms indicate the distribution of overlap (Jaccard) scores for the given dataset, and the labelled dashed line indicates the mean of this distribution. The individual modules shown here as well as for the KEGG Pathways and IntAct Complexes module standards can be found in **Supplementary Data File 4**. The LSM2-7 complex is indicated by the arrows. PPI = protein-protein interaction. This analysis uses the same networks and BiONIC features as **Fig. 2**.

**Figure S5.**
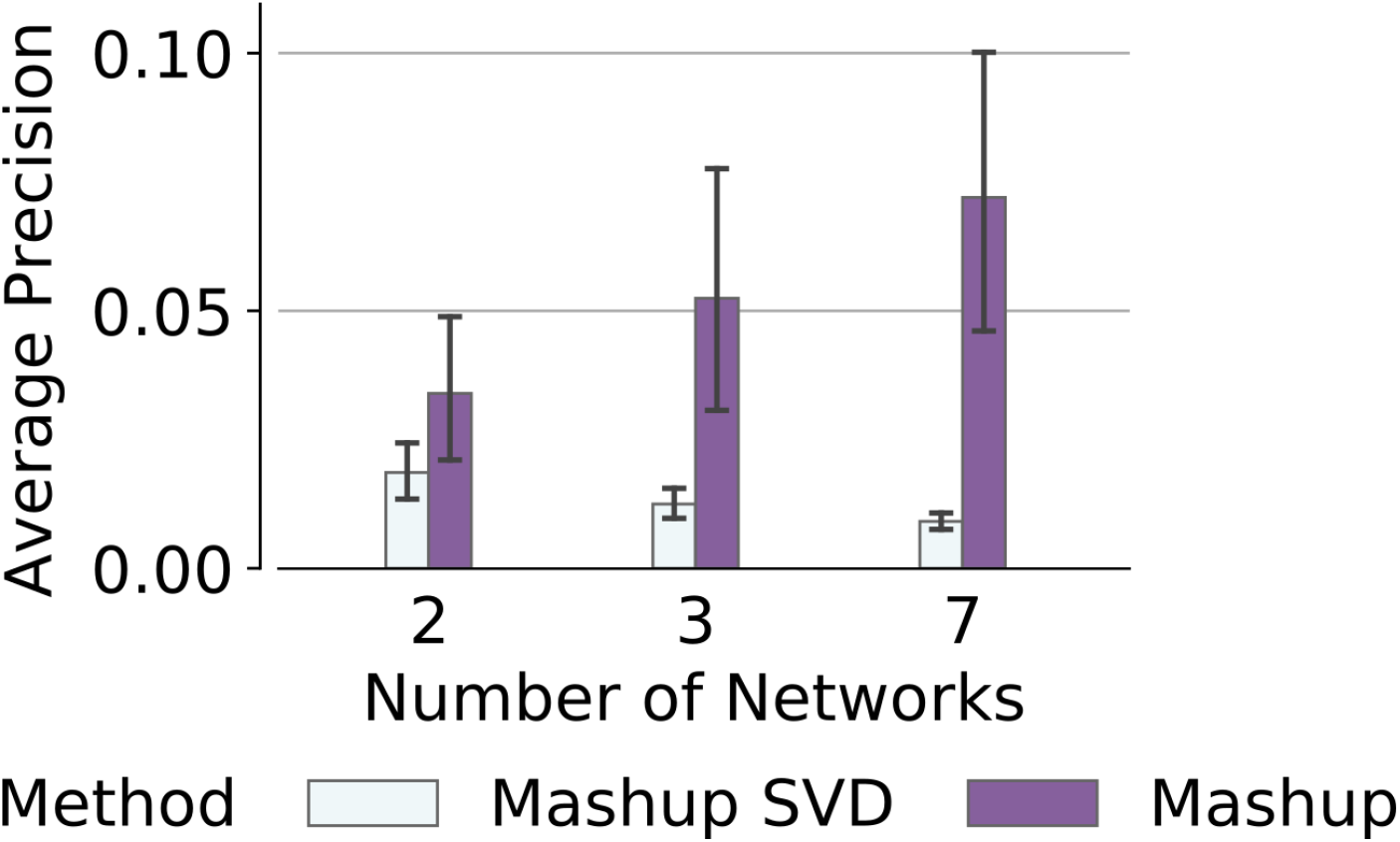
Comparison of Mashup with the author provided singular value decomposition approximation of Mashup (denoted Mashup SVD). Random sets of yeast co-expression networks were sampled, integrated and scored against the KEGG pathways co-annotation standard, analogous to **Fig. 4a**. Error bars indicate the 95% confidence interval.

**Figure S6.**
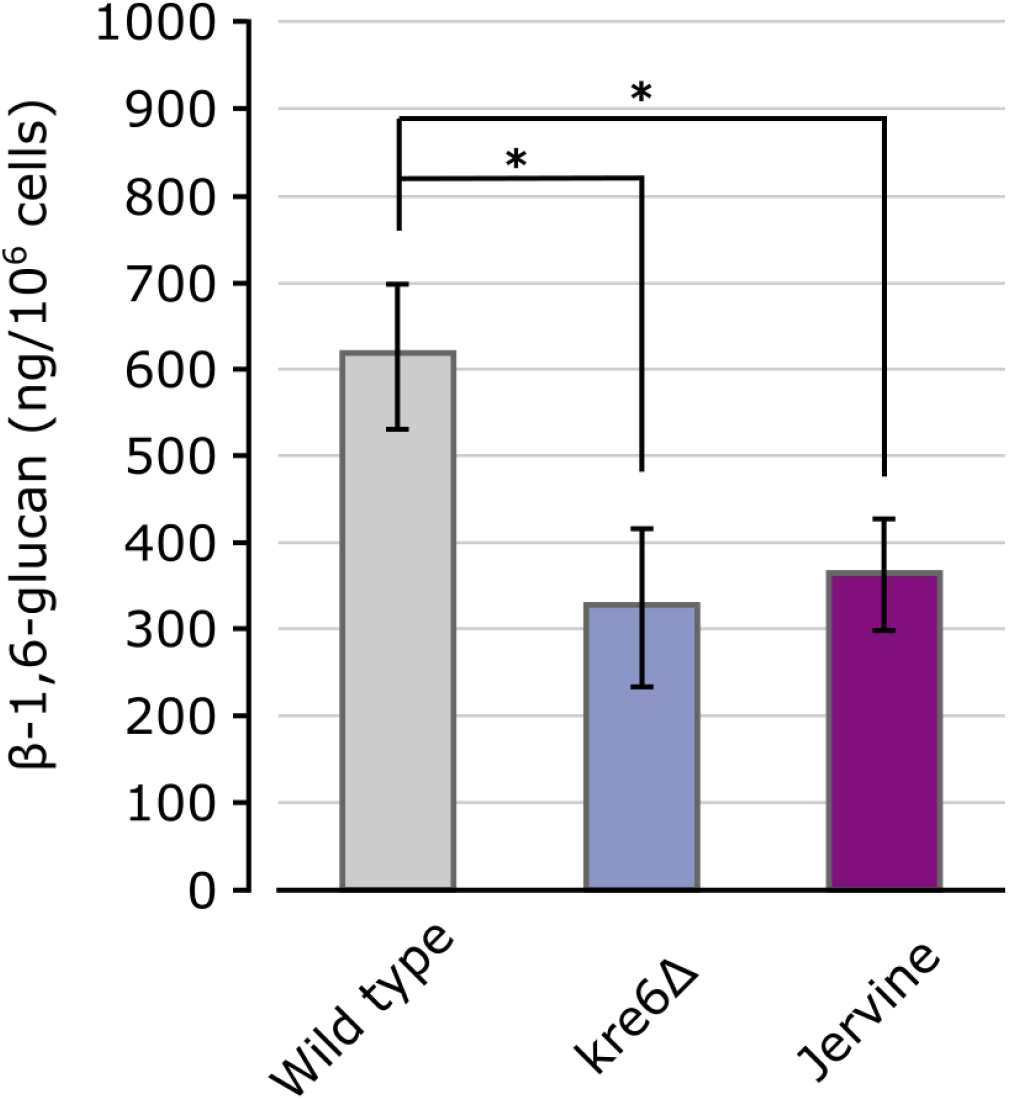
β-1,6-glucan levels in yeast strains. The amount of glucan per cell was calculated using pustulan as a standard. * Significant difference (p < 0.05 after Bonferroni correction, t-test).

## Supplementary Data Files

[Supplementary Data File 1](https://figshare.com/articles/dataset/Hyperparameter_Optimization_Results/15025995)

**Hyperparameter Optimization Results**. Hyperparameter optimization results across integration methods integrating three *Schizosaccharomyces pombe* networks. The chosen (best) hyperparameter combinations for each method are highlighted.

[Supplementary Data File 2](https://figshare.com/articles/dataset/Integrated_Networks_Details_xlsx/14866917)

**Integrated Network Details**. Publication, gene count, edge count and experimental type for each yeast network and each human network used in **Fig. 2-6**. Rows in yellow indicate the three yeast networks used in **Fig. 2-4, 6** integrations.

[Supplementary Data File 3](https://figshare.com/articles/dataset/Figure_2_Evaluation_Standards_Details/14983683)

**Evaluation Standards Details**. Gene count, co-annotation count, module count and class count details for each standard used in the **Fig. 2-5** evaluations.

[Supplementary Data File 4](https://figshare.com/articles/dataset/Figure_2_Module_Detection_Results/14975298)

**Module Detection Results**. Overlap of standard-optimized clusters obtained from the **Fig. 2c** module detection analysis for networks as well as integration methods. Module standards are IntAct Complexes, KEGG Pathways and GO Biological Processes.

[Supplementary Data File 5](https://figshare.com/articles/dataset/Fig_S3_S4_Module_Detection_Results/14976747)

**Fig. S3, S4 Module Detection Results**. Overlap of known per-module-optimized clusters obtained from the **Fig. 2, 3** input networks and integration methods, with IntAct Complex, KEGG Pathway and GO Biological Process modules.

[Supplementary Data File 6](https://figshare.com/articles/dataset/50_Compound_TS_Allele_Screen_Results/16645894)

**50 Compound TS Allele Screen Results**. Files containing the TS allele CG-scores and IQR-scores, screened against 50 compounds (at multiple concentrations) that were selected by BIONIC.

[Supplementary Data File 7](https://figshare.com/articles/dataset/Essential_Gene_Compound_Sensitivity_Predictions/15158037)

**Essential Gene Compound Sensitivity Predictions**. Essential yeast gene compound sensitivity predictions for 50 selected compounds using BIONIC.

[Supplementary Data File 8](https://figshare.com/articles/dataset/BIONIC_Yeast_Features/16632286)

**Integrated BIONIC Features**. Learned BIONIC features from yeast networks (protein-protein interaction, co-expression, and genetic interaction) integrated and used in **Fig. 2, 3, 6**.

[Supplementary Data File 9](https://figshare.com/articles/dataset/Evaluation_Standards/16629139)

**Evaluation Standards**. Yeast evaluation standards for co-annotation prediction, module detection, and gene function prediction used in **Fig. 2a, 3a, 4, 5a** as well as the human co-annotation standard used in **Fig. 5b**.

